# Punctuated Evolution of Endomembrane Compartments in Proto-Eukaryotes

**DOI:** 10.64898/2026.04.13.718263

**Authors:** Sahana Shridhar, Kritika Kumari, Mukund Thattai

## Abstract

Eukaryotic cells are defined by their endomembranes: compartments such as the endoplasmic reticulum (ER), Golgi and endosomes, exchanging cargo via vesicles. The evolutionary origins of endomembrane compartments remain unclear. Here we construct molecular-evolutionary trajectories for the stepwise addition of compartments after the emergence of the proto-ER in an ancestral eukaryote. We represent compartments and vesicles as nodes and edges of a directed graph. Vesicle budding and fusion regulators such as coats and SNAREs control cargo flows and determine compartment compositions. We computationally sample billions of possible graphs, and enumerate how duplication, deletion and mutation of regulators drive graph transitions. We find that evolutionary trajectories display punctuated shifts in compartment composition and number, interspersed with thousands of neutral mutations. The first added compartment inherits functions from the proto-ER or plasma membrane, or gains novel functions. Our results show how, given a billion years, simple molecular steps can generate complex endomembrane systems.

**Significance Statement:** Eukaryotic cells contain a system of endomembrane compartments that sort, process and deliver molecules to precise cellular destinations. This endomembrane system is a defining feature of all complex life, yet its evolutionary origins remain obscure. How did a proto-eukaryote with a single ancestral endomembrane compartment evolve into a cell with a Golgi, endosomes, lysosomes and other compartments characteristic of modern eukaryotes? We model this process from first principles, connecting the duplication, deletion and mutation of molecular regulators to compartment gain or loss. We find a punctuated pattern of endomembrane elaboration: a long phase of neutral exploration, driven by the mutation of duplicate gene copies, precedes the emergence of new compartments and functions.

## I. Introduction

The defining features of the eukaryotic cell plan are the product of a billion years of evolution. Mitochondria, the nucleus, and the endomembrane system of compartments exchanging vesicular cargo emerged in successive steps whose timing and mechanism are active areas of research [1–3]. The First Eukaryotic Common Ancestor (FECA) diverged from the Asgard archaeal lineage prior to the Great Oxidation Event, over 2.4 billion years ago [4, 5]. FECA was a prokaryote that possessed certain eukaryote-like traits [2, 3, 6] but likely lacked stable endomembrane compartments. The Last Eukaryotic Common Ancestor (LECA) dated to between 1.8 and 1.2 billion years ago [4, 7]. LECA possessed symbiotic mitochondria alongside an archaea-derived endomembrane system of interconnected compartments [8, 9]. The modern eukaryotic cell plan emerged through a series of steps on the path from FECA and LECA.

Three key features likely arose relatively early in eukaryogenesis, in an unknown order [3, 10, 11]: A bacterial endosymbiont was acquired within the archaeal host, giving rise to mitochondria [12, 13]. The proto-endoplasmic reticulum (pER) emerged as a stable endomembrane compartment, part of which formed the nuclear envelope [11]. And transport vesicles arose to ferry lipids and membrane-integral proteins between endomembranes and the plasma membrane (PM) [3]. These three features define a rudimentary proto-eukaryotic endomembrane system. We are interested in how new intracellular compartments (ICs) such as the Golgi apparatus, endosomes and lysosomes were layered onto this ancestral state [14– The budding and fusion of transport vesicles from endomembrane compartments is regulated by conserved molecular modules [17, 18]. Coat/adaptor complexes select specific cargo and drive vesicle budding from source compartments [19, 20]. SNARE proteins mediate the fusion of vesicles to specific target compartments [21, 22]. These regulators control how vesicular cargo flows between compartments, and therefore determine compartment compositions [23– Proto-eukaryotes inherited a minimal molecular toolkit for vesicle traffic from their Asgard archaeal ancestors [3, 26–2 Gene duplication, deletion and mutation of these regulators over the next billion years generated the genomic complexity seen in LECA [6, 15, 16, 30]. Paralogous regulator copies are often localized to distinct endomembrane organelles [31]. The organelle paralogy hypothesis (OPH) connects these genomic and cell-biological observations: it posits that the duplication of regulatory modules paralleled the emergence of new compartments [32– Mathematical models have shown that duplicate copies of regulatory molecules support the co-existence of multiple compartment types [23– But these models do not incorporate evolutionary dynamics: they do not show how molecular evolution can drive the emergence of new compartments given a pre-existing endomembrane system.

Here we ask: starting from a proto-eukaryote with a pER, a PM, and a minimal vesicle traffic toolkit, what evolutionary paths lead to the emergence of new endomembrane compartments? We address this question with a computational model that includes dynamics at multiple levels: genomic (gene duplication, deletion, mutation); molecular (budding and fusion modules); and cellular (compartments and vesicle flows). Compartments and vesicles are represented as the nodes and edges of a graph. Molecular regulators determine which graphs are physically allowed, and molecular changes drive transitions between allowed graphs. We enumerate and assign a fitness consequence to each transition, and construct all viable evolutionary trajectories from the proto-eukaryotic starting point. These trajectories reveal a striking punctuated pattern of endomembrane evolution [35–37]: compartment gains require long periods of neutral exploration, enabled by the availability of redundant or duplicate genes. Our results suggest a complex connection between genotypic and phenotypic changes, and highlight how contingency, canalization and exaptation shape endomembrane evolution.

## Results

### Molecular regulators of vesicle budding and fusion

We use “compartments” to refer to large stable membrane-bounded structures, and “vesicles” to refer to smaller transient membrane-bounded carriers that mediate cargo flows between compartments. Vesicle budding and fusion are regulated by modules of interacting proteins [14, 17]. We assume that specific membrane-integral molecules ultimately determine the properties of compartments and vesicles, though these processes may make use of shared cytoplasmic factors.

Coat/adaptor complexes (henceforth referred to as coats) such as clathrin-AP, COPI or COPII [19, 20] select cargo and drive vesicle budding at source compartments. These cytoplasmic complexes are recruited by specific compartmental markers through a reaction cascade involving Arf GTPases [38]. Coats and their recruiters comprise the budding module. Once a vesicle is formed at a source compartment, we assume it samples all possible target compartments by diffusion or cytoplasmic mixing [39]. Specific fusion occurs when v-SNAREs on the vesicle bind to cognate t-SNAREs on the target compartment [21, 22]. SNAREs operate in conjunction with Rab GTPases [40, 41], tethering complexes [42, 43] and other regulators [18, 21, 44, 45]. v-SNAREs that actively drive the fusion of one vesicle are recycled as inactive cargo on other vesicles [24]. We consider three models of SNARE regulation (Fig. S1E,F). In the activator model, a v-SNARE is active only when its activator is present on the same vesicle; in the inhibitor model, a v-SNARE is active except when its inhibitor is present on the same vesicle; in the unregulated model, a v-SNARE is always active. SNAREs and their regulators comprise the fusion module.

### Cargo flow in the compartment graph

We represent the endomembrane system as a compartment graph: a directed graph whose nodes are compartment types and whose edges are vesicle flows between compartments (Methods, Section A; Fig. 1B,C). We assume compartments with identical compositions have identical properties, so we collapse multiple instances of each compartment type into a single node. The simplest proto-eukaryotic endomembrane system contains two compartments. First, the proto-endoplasmic reticulum (pER): This is defined as the compartment contiguous with the nuclear envelope, where all membrane-integral cargo are synthesized [46]. Second, the plasma membrane (PM): This is the interface between the cell and its environment. Note that the ancestral pER and PM may host functions typically associated with other compartments in modern eukaryotes. Here we explore the addition of new intracellular compartments (IC) to the proto-eukaryotic initial state, considering graphs with up to four compartments (pER, PM, IC1, IC2).

**Fig. 1:**
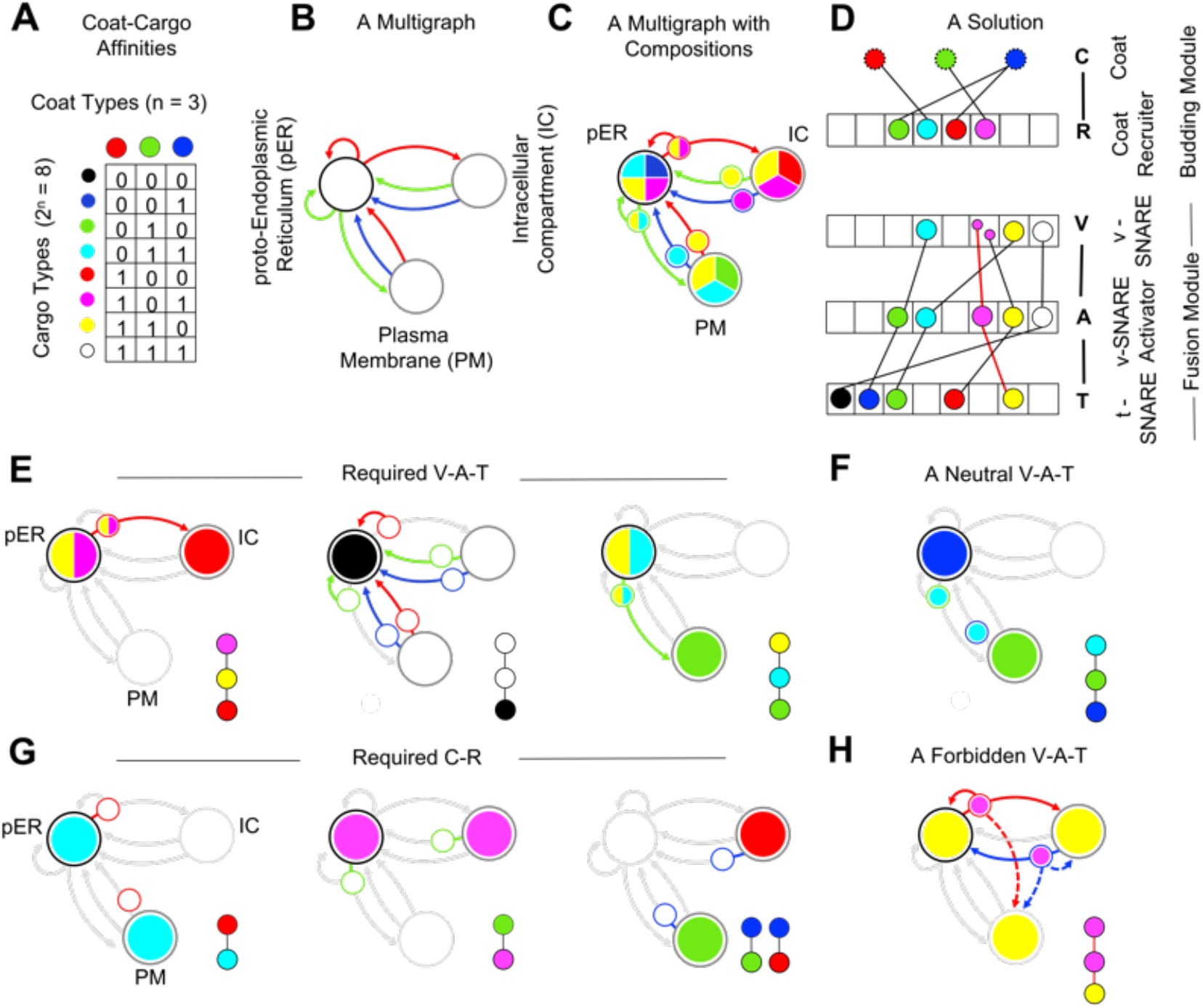
Emergence of the compartment graph from molecular interaction rules. **(A)** Coat-cargo affinities. Each cargo type binds to a specific subset of coat types. This determines which vesicles it is loaded into and which compartments it can reach. For *n* =3 coat types (edge colors) there are 2^*n*^ =8 cargo types (binary numbers 0 to 7, cargo colors). Cargo type 0 (black, only on the pER) and cargo type 7 (white, on every compartment) are not explicitly shown. **(B)** A compartment multigraph (a directed graph including self-edges and multiple edges between nodes) is constructed by assigning coat labels to edges, and pER, PM or IC identities to nodes. Each labeled multigraph represents a candidate endomembrane topology. **(C)** Coat labels on edges, together with the pER as the source of cargo synthesis, uniquely determine the composition of every compartment and vesicle (Fig. S1A,B). We represent compositions by cargo-colored wedges on compartments (large circular nodes) and vesicles (small circles on edges). **(D)** We say a graph is allowed if it has at least one molecular solution, obtained by selecting budding and fusion modules that enable all present vesicle flows without enabling any absent ones. Each budding module is a doublet (a single edge) comprising a coat (C) and its specific recruiter (R). Each fusion module is a triplet (a pair of linked edges) comprising a v-SNARE (V), its specific regulator (activator A or inhibitor I), and its cognate t-SNARE (T). Each component of a budding or fusion module is assigned to one of the 2^*n*^ cargo types (colors). Here we show a molecular solution for the activator model, see Fig. S1E,F for other SNARE regulation models. Each required module (black edges) can be linked to some vesicle budding or fusion event present in the graph. We also show a forbidden fusion module (red edges) which would lead to a vesicle fusion event absent from the graph. **(E)** Required fusion modules (V-A-T triplets) and the vesicle fusion events they enable. **(F)** A neutral V-A-T module that has no effect on the graph since its components never co-localize on the same vesicle. **(G)** Required budding modules (C-R doublets) and the vesicle budding events they enable. **(H)** A forbidden V-A-T module (red edges in panel D) that would enable a vesicle flow absent from the graph (dotted red edge in the graph). See Fig. S1 for details on cargo flows and SNARE regulatory models.

We assume there are *n* available coat types. Each membrane-integral molecule is assigned a cargo type, defined by the set of coat types it can bind to (Fig. 1A). Two molecules of the same cargo type behave identically, flowing between the same compartments on the same vesicles regardless of their identity. This abstraction collapses the enormous biochemical diversity of the endomembrane system onto a small set of 2^*n*^ transport behaviors. Every cargo type starts at the pER and flows along any vesicle with a compatible coat. Cargo with no coat affinity remains confined to the pER; cargo with affinity for all coats reaches every compartment. Any intermediate cargo type eventually reaches a subset of connected compartments within which it cycles (Fig. S1A,B). The degradation of cargo is balanced by replenishment from the pER. We assume degradation is slow compared to the transport rate [47, 48]. Cargo will therefore be present in high amounts only on cyclic routes, and in trace amounts on unidirectional replenishment routes (Fig. S1A,B). This completely defines the cargo types found on every compartment and vesicle (Methods, Section B; Fig. 1C). We represent this information as a multigraph: a directed graph allowing self-edges and multiple edges between nodes. The fully annotated multigraph has coat labels on edges, and cargo compositions assigned to compartments and vesicles (Fig. 1B,C).

### Molecular solutions of compartment graphs

Every compartment graph must satisfy a self-consistency condition. We have seen how cargo flows determine compartment and vesicle compositions. These compositions must, in turn, enable the cargo flows. Specifically, every vesicle flow that is present in the graph must have a budding module at its source and a fusion module at its target, and no module may inadvertently enable a vesicle flow that is absent.

Budding and fusion modules are made up of several membrane-integral regulatory molecules: coat recruiters, v-SNAREs, v-SNARE activators or inhibitors, and t-SNAREs (Methods, Section C; Fig. 1D). Multiple paralogous copies of these modules can simultaneously exist. Each membrane-integral component of each regulatory module is assigned to one of 2^*n*^ cargo types. A budding module is a coat-recruiter (C-R) doublet: one of *n* coat types paired with one of 2^*n*^ recruiter cargo types, giving *n·*2^*n*^ possibilities. The fusion module in the activator model is a v-SNARE–activator–t-SNARE (V-A-T) triplet: each component is drawn from among 2^*n*^ cargo types, giving 2^3*n*^ possibilities. In the 3-coat activator model this yields 24 possible budding modules and 512 possible fusion modules.

The genome encodes a specific selection of modules from this space of possibilities, and must be checked against the self-consistency conditions of the compartment graph. A set of budding and fusion modules satisfying these conditions is a molecular solution (Methods, Section D; Fig. 1D-H). A compartment graph that has at least one molecular solution is an allowed graph. A single graph may have multiple molecular solutions; a single solution may be compatible with multiple graphs. We computationally sample billions of possible compartment graphs, spanning 2 to 4 compartments and 2 to 3 coat types (Fig. S1C,D). We find that allowed graphs are extremely rare, and their fraction decreases sharply as the number of compartments and vesicle flows increases (Table I). This observation motivates the central question of our study: How could new compartments arise via a molecular-evolutionary process, given that physically allowed graphs are so rare?

**TABLE I:** Allowed compartment graphs across SNARE regulatory models. Graph nodes represent compartments such as the proto-endoplasmic reticulum (pER, where all membrane-integral cargo are synthesized), the plasma membrane (PM, considered as a compartment whose “interior” is the extracellular space), or other intracellular compartments (IC). Graph edges represent vesicle flows, where each vesicle type is associated with one of 2 or 3 types of cargo-selective coats. We test all possible 2-compartment (pER/PM), 3-compartment (pER/PM/IC), and 4-compartment (pER/PM/IC1/IC2) graphs, up to some maximum number of edges (Fig. S1C,D). (Going to higher coat and compartment numbers is computationally infeasible due to the combinatorial explosion of possible graphs.) For each possible graph we search for a molecular solution under the activator, inhibitor, and unregulated SNARE regulatory models.

| Coat Types | Cmpts/<br>Edges | Graphs<br>Tested | Allowed<br>Activ. | Allowed<br>Inhib. | Allowed<br>Unreg. |
| --- | --- | --- | --- | --- | --- |
| 2 | 2/8 | 90 | 13 | 23 | 11 |
|  | 3/12 | 92,022 | 26 | 91 | 16 |
| 3 | 2/12 | 3,622 | 235 | 624 | 117 |
|  | 3/17 | 75,954,566 | 6,990 | 92,910 | 1,626 |
|  | 4/14 | 25,411,159,020 | 168 | 1,266 | 0 |

### Required, forbidden and neutral molecular modules

Given a multigraph with coat labels and compositions, the set of possible modules splits into three classes, defined by the graph condition (Methods, Section E; Fig. 2A). There is one set of required modules associated with each vesicle budding and fusion event present in the graph; if multiple modules satisfy the same event, having more than one is redundant. There is a set of forbidden modules whose presence would enable budding and fusion events absent from the graph. Neutral modules do not affect the graph. Every molecular solution involves selecting at least one module from each required set, while having no modules in the forbidden set (Fig. 1D-F). An impossible graph is one for which some set of required modules falls completely within the set of forbidden modules (Fig. 2B). The probability that a graph is impossible increases with the number of compartments and vesicle flows, explaining the rarity of allowed graphs in Table I.

**Fig. 2:**
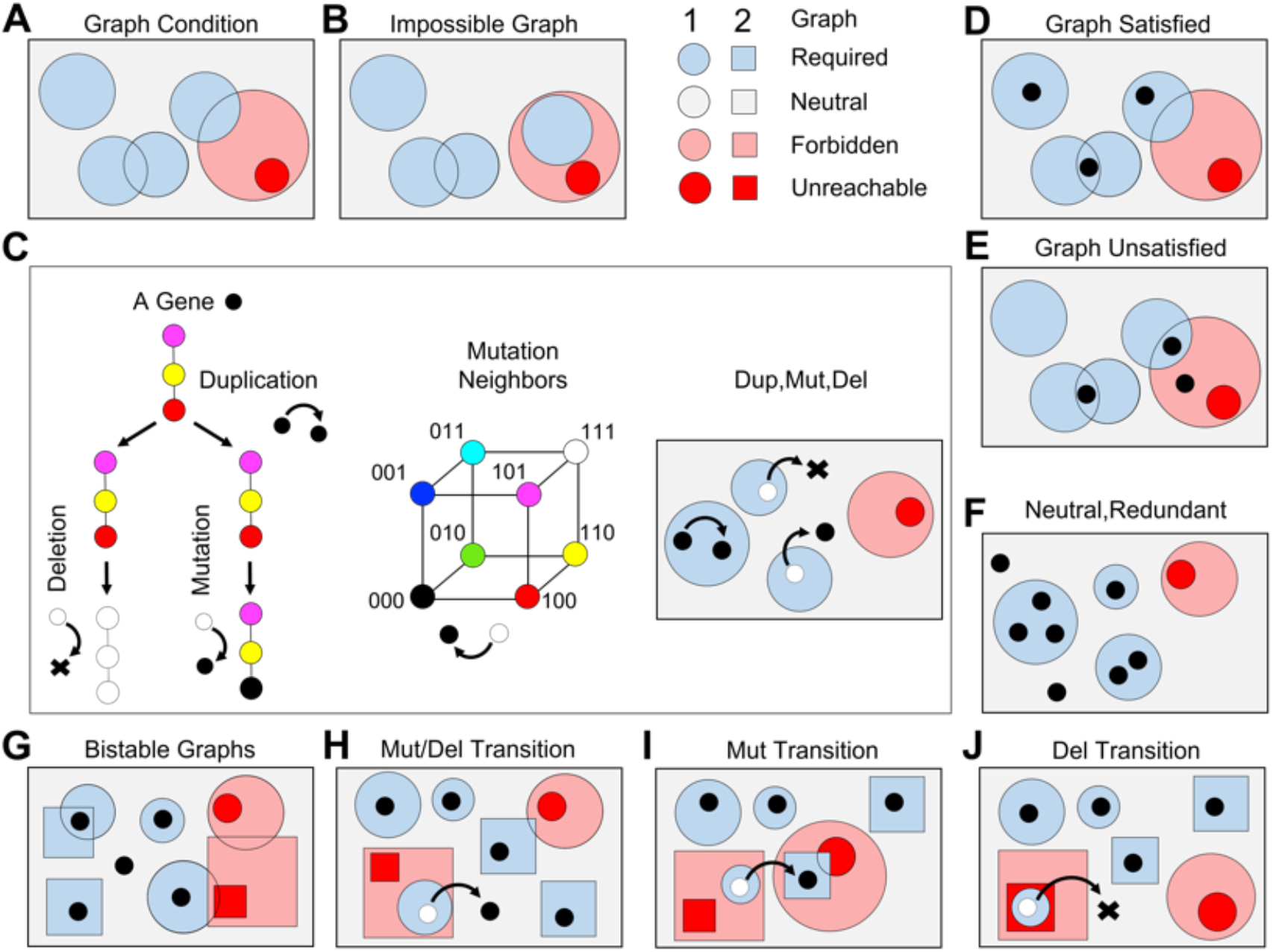
Graph transitions by elementary evolutionary moves. We are given an initial graph (1, circles) and a final graph (2, squares). We ask whether every molecular solution of the initial graph can be converted to some molecular solution of the final graph using only elementary evolutionary moves. **(A)** The graph condition. The rectangle represents the space of all possible budding and fusion modules. For the 3-coat system there are 24 possible C-R budding modules and 512 possible V-A-T fusion modules (Fig. S2A-D). Each blue region shows modules that are required to enable each vesicle budding and fusion event present in a graph. The pink region shows modules that are forbidden because they would enable vesicle budding and fusion events absent in a graph. The gray region shows neutral modules that have no effect on the graph. The red region shows forbidden modules that are unreachable by mutations from the neutral region. **(B)** An impossible graph. One of the required regions is completely contained within the forbidden region, so we cannot find a molecular solution. **(C)** The three elementary evolutionary moves. Black dots represent modules (C-R doublets or V-A-T triplets) that are actually encoded by the genome. Duplication adds a new copy of an existing module. Deletion removes a module. Mutation flips a single coat affinity, moving a cargo type of a module component to a neighboring vertex in coat-affinity space. **(D)** The graph condition is satisfied when there is at least one module within each required region and there are no modules within the forbidden region. **(E)** The graph condition is unsatisfied if some required region contains no modules, or the forbidden region contains a module. Modules outside the required and forbidden regions are neutral. Extra modules within any required region are redundant. **(G)-(J)** Graph transition types. We consider special molecular solutions of graph-1 that are a single elementary move from a molecular solution of graph-2. This figure is schematic, but we have explicit representations of required, forbidden and neutral regions for every graph in our dataset. **(G)** Bistable transition. Both graph-1 and graph-2 conditions can be simultaneously satisfied, the initial condition selects one of them. **(H)** Mutation/deletion transition. A required region of graph-1 falls in the forbidden region of graph-2. A single mutation into the neutral region, or a single deletion, un-satisfies the graph-1 condition and satisfies the graph-2 condition. **(I)** Mutation-only transition. A required region of graph-1 falls in the forbidden region of graph-2, and vice versa. A single mutation is required to un-satisfy the graph-1 condition and satisfy the graph-2 condition. **(J)** Deletion-only transition. A required region of graph-1 falls in the unreachable forbidden region of graph-2. A single deletion is required to un-satisfy the graph-1 condition and satisfy the graph-2 condition. See Fig. S2 for statistics of required and forbidden regions.

### Elementary evolutionary moves and graph transitions

Starting with a molecular solution, we define three elementary evolutionary moves (Methods, Section F; Fig. 2C). Duplication adds a copy of an existing module. Deletion removes an existing module. Mutation flips a single coat affinity of a single module component, akin to modifying the coat-selective cytoplasmic tail of a transmembrane protein [49]. Mutation can shift a module between required, forbidden and neutral classes. The duplication, deletion and mutation of neutral and redundant modules allow a large genomic space to be explored without impacting the structure of a graph.

Suppose we want to determine if a given initial graph-1 can evolve into a given final graph-2 (Methods, Section G). We must check if every solution of graph-1 can be transformed into some solution of graph-2 using only elementary evolutionary moves. Each step must therefore be neutral with respect to graph-1, until we reach a solution of graph-2. This structure, of a neutral walk followed by a sudden transition, is central to our analysis. The critical step is the final one, which precipitates one of four possible types of graph transitions (Fig. 2G-J; Fig. S2E). In bistable transitions the conditions of both graphs can be satisfied by the same molecular solution, so the initial condition selects one of the graphs. In the remaining transition types the final step forces a shift from graph-1 to graph-2. In mutation/deletion transitions a single mutation or deletion removes a forbidden module of graph-2. In mutation-only transitions a single mutation simultaneously removes a forbidden module and adds a required module of graph-2. In deletion-only transitions a forbidden module of graph-2 cannot be removed by mutation alone, and must be eliminated by deletion. We can explicitly determine if a neutral walk exists from a graph-1 solution to a graph-2 solution, based purely on the two graph conditions.

### Affordance framework for endomembrane function

Endomembrane compartments offer many potential benefits to the cell, including enhanced nutrient uptake, biosynthetic control, and the localization of incompatible processes to distinct organelles [50– Graph transitions change the structure of the endomembrane system, as compartments are gained or lost and vesicle flows are rewired. A transition that destroys an existing cellular function would be strongly selected against. The challenge in modeling this is that we do not know the function of any given compartment graph, so we cannot quantify how it contributes to cellular fitness.

We resolve this by using the concept of affordance [54], a general framework for understanding how objects offer opportunities for action (Methods, Section H). We say a compartment carries an affordance label if at least one cargo type is found exclusively on that compartment, making it molecularly distinguishable from all others. Similarly, a vesicle flow carries an affordance label if it connects two labeled compartments, and at least one cargo type from the source compartment travels exclusively along that flow. The set of compartments and vesicle flows with affordance labels collectively define the graph’s affordance motif. This motif does not specify a function. Rather, it shows the potential of a graph to support any function. A unique cargo that labels a compartment or vesicle provides a simple foundation upon which to build a function, when compared to a more complex combinatorial code. Each label may be associated with one or more functions. Affordance labels allow cells to robustly target distinct molecules to distinct compartments and vesicles, and thereby derive a fitness benefit.

### The affordance ratchet and evolutionary trajectories

A compartment graph can be represented at several levels of detail (Fig. 3A,B). The multigraph representation is the most detailed: it shows coat labels on edges and cargo compositions on every compartment and vesicle. The simple graph representation is coarse-grained: it retains compartments with cellular identities (pER, PM, IC), discards self-edges and multi-edges, and discards coat labels and compositions. The affordance label representation explicitly shows which cargo types act as compartment or vesicle labels. The affordance motif is the coarsest representation: it retains only the affordance-labeled compartments and flows, discarding everything else.

**Fig. 3:**
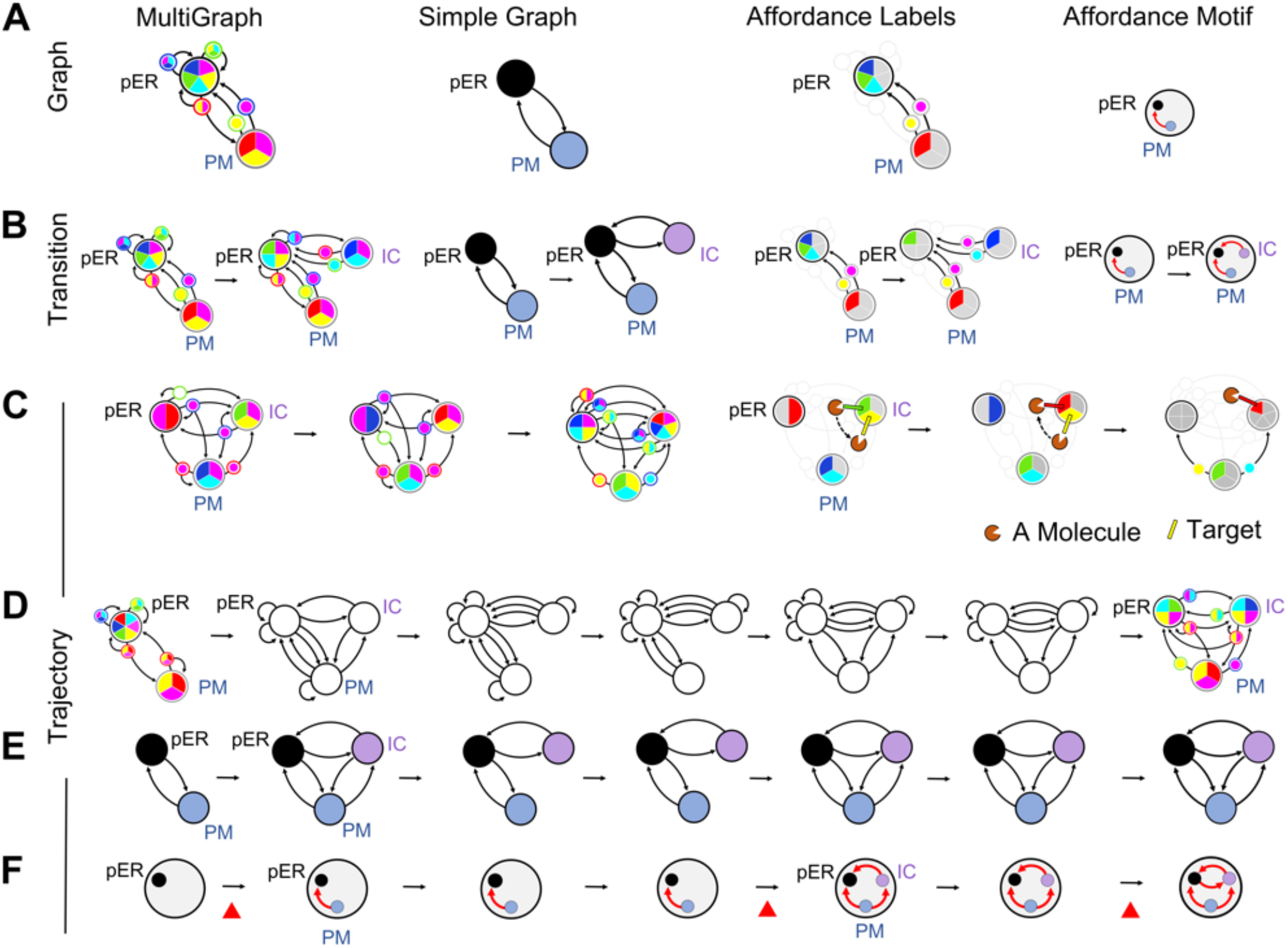
Affordance motifs and evolutionary trajectories. **(A)** An allowed graph from the 3-coat activator model shown in multiple representations. A compartment is said to have an affordance label if there is at least one cargo type that is found exclusively on that compartment. A vesicle flow is said to have an affordance label if it connects two labeled compartments, and some cargo type from the source compartment flows exclusively to the target compartment (not back to the source compartment along self-edges, nor to some third compartment). The multigraph representation shows coat labels on edges and cargo compositions on every compartment and vesicle. The simple graph representation shows compartment identity (pER, black; PM, blue; IC, purple) and vesicle flows, discarding self-edges and multi-edges. The affordance label representation shows cargo types that exclusively label compartments and vesicle flows. The affordance motif representation shows only those compartments and vesicle flows that have affordance labels, capturing the functional potential of the compartment graph. **(B)** We say a graph transition is allowed if it can be achieved by elementary evolutionary steps, and the affordance motif is either preserved or expanded. This defines the “affordance ratchet”. We show an allowed graph transition in all four representations (Fig. S2A,B). **(C)** Example of the affordance motif remaining unchanged even when the specific labels turn over. A functional molecule is first localized at the IC compartment by binding the green label. A shift from the green to the yellow label is neutral. A graph transition removes the green label but adds a red label. A shift from the yellow label to the red label is neutral. A second graph transition removes the yellow label. The protein is correctly localized throughout this process. **(D-F)** An evolutionary trajectory, defined as a sequence of graph transitions consistent with the affordance ratchet. Each step (rightward arrow) involves a neutral walk (see Fig. 5E). **(D**,**E)** At the multigraph and simple graph level, vesicle flows and compartments are gained or lost at each transition. **(F)** At the affordance motif level, affordances are preserved or expanded (red triangle) at each transition. The trajectory shown here starts with the simplest proto-eukaryotic compartment graph with only the labeled pER and an unlabeled PM. It ends with a compartment graph from which no further affordance-expanding transitions are possible. See Fig. S3 for more examples of transitions.

A graph’s affordance motif may be preserved even if the specific cargo types that act as labels turn over (Fig. 3C). These various representations allow us to track transitions at different levels of abstraction: the multigraph changes at every graph transition, the simple graph and affordance labels change less frequently, and the affordance motif changes least of all. We can stitch transitions into complete trajectories, while imposing an affordance ratchet (Methods, Section H; Fig. 3B,C; Fig. S3A,B). Transitions that destroy an existing affordance motif are selected against. Transitions that preserve the affordance motif are treated as functionally neutral, and are reversible. Transitions that add new affordances while preserving existing ones represent evolutionary leaps, and are irreversible: they expand the functional repertoire of the endomembrane system. Every trajectory begins with the proto-eukaryotic initial graph, and terminates at an endpoint graph from which no affordance-expanding transition exists. Compartments and vesicle flows may be gained or lost along the way, provided the affordance motif is preserved or expanded.

### Contingency in evolutionary trajectories

We can compare how connected the evolutionary landscape is, across SNARE-regulatory models. We have already explicitly enumerated all allowed compartment graphs for the 3-coat activator, inhibitor and unregulated models (Table I). We now enumerate all allowed graph transitions consistent with elementary evolutionary moves and the affordance ratchet. We can represent this as an adjacency matrix whose entries are 1 if a directed transition is allowed, and 0 otherwise. For *n*_*g*_ graphs there are 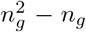 possible transitions. If *n*_*t*_ transitions are allowed, the density of the adjacency matrix is 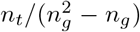. The unregulated model has 1,743 graphs and 105,033 transitions, giving a density of 3.5%. The activator model has 7,393 graphs and 1,430,605 transitions, giving a density of 2.5%. The inhibitor model has 94,800 graphs but too many transitions to practically enumerate; sampling gives a density of approximately 4.1% (10,000 allowed transitions from 245,323 samples).

We simulate evolutionary trajectories by Monte Carlo sampling, starting from the proto-eukaryotic initial graph (Methods, Section I). At each step we jump to a randomly chosen neighbor of the current graph, as allowed by the adjacency matrix. (This approach can be extended with a weighted probability across neighbors, and with occasional transitions that violate the affordance ratchet; these extensions do not significantly change our results.) We find strong statistical regularities in these trajectories. Certain simple graphs and affordance motifs never occur in the set of allowed graphs (Fig. 4A-C), reflecting the impact of molecular constraints [24]. For the 3-coat activator model all trajectories terminate at one of nine endpoint affordance motifs (Fig. 4B). The set of trajectories converging at each endpoint are diverse, involving multiple changes in graph topology through compartment and vesicle gains and losses (Fig. 4D). Strikingly, the symmetry or asymmetry of these endpoints with respect to the pER, PM and IC labels is already reflected in the early steps of each trajectory, revealing how historical contingency restricts later evolutionary possibilities (Fig. 4D; Fig. S4).

**Fig. 4:**
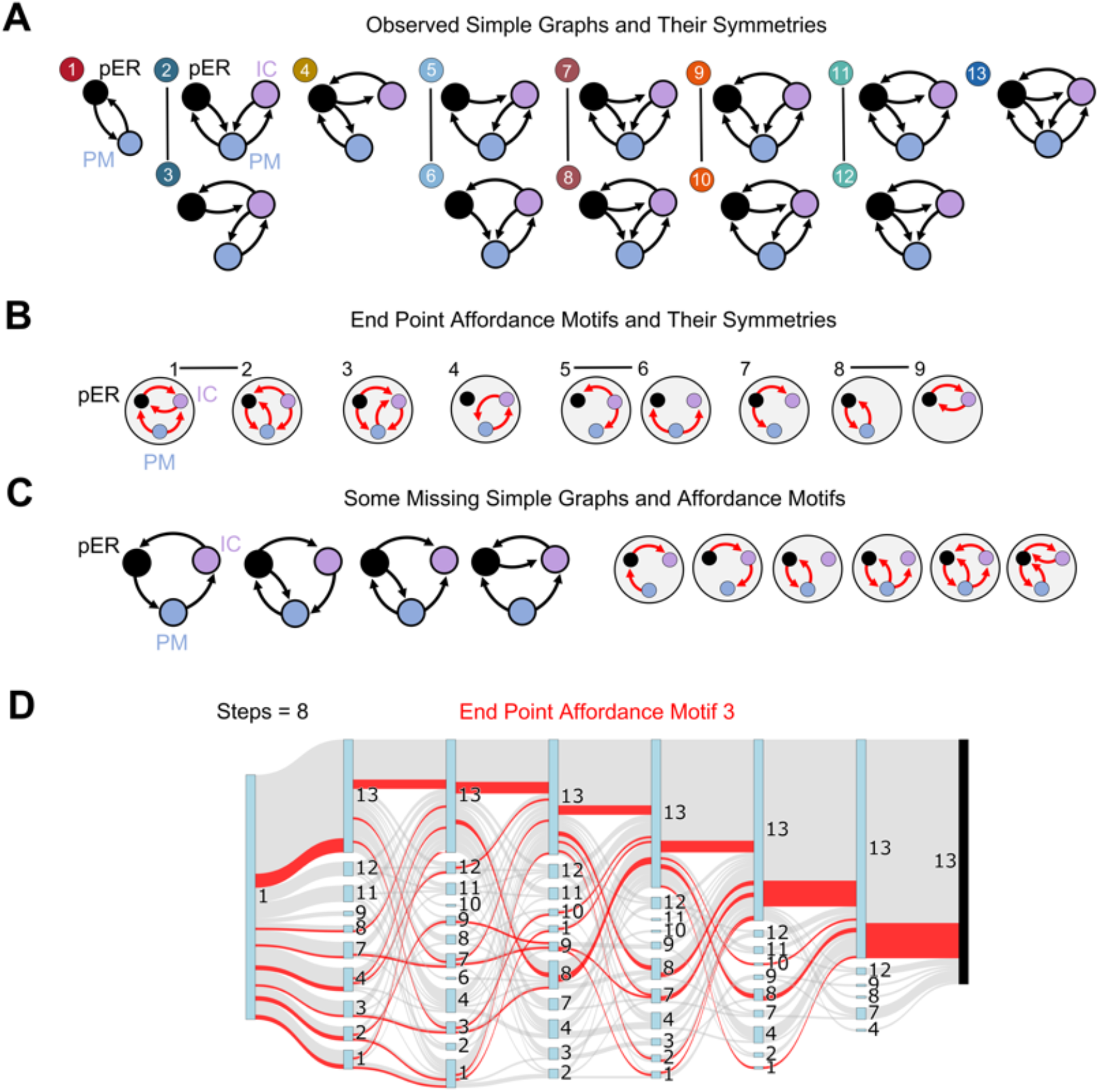
Patterns of observed compartment graphs and trajectories. **(A)** The full set of simple graphs observed in the 3-coat activator model. We highlight symmetry pairs: graphs that correspond by exchanging the PM and IC labels. **(B)** The nine endpoint affordance motifs reached by the complete set of sampled trajectories. Lines link PM IC symmetry pairs. **(C)** Examples of simple graphs and affordance motifs that are topologically possible, but which correspond to impossible graphs. This reflects the filtering effect of molecular constraints. **(D)** A Sankey diagram illustrating historical contingency. We show 188 simulated trajectories that start from the proto-eukaryotic compartment graph and terminate within 8 steps (inclusive of the first). Those trajectories that terminate at a graph with affordance motif 3 are highlighted in red. Each trajectory goes through a different history of simple graph types, numbered as in (A). Early events constrain later ones, so the diversity of simple graphs narrows near the endpoint. For example, trajectories passing through simple graph type 11 at any stage cannot reach endpoint affordance motif 3. See Fig. S4 for more details on endpoint-specific patterns.

### Canalized landscape of endomembrane evolution

We next explore the global structure of the trajectories between the proto-eukaryotic starting point and the endpoints. The allowed compartment graphs and their transitions define a large space involving thousands of graphs. This space has a natural hierarchical structure (Fig. 5A). At the finest scale, individual graphs connect to each other through elementary transitions. We group these graphs into classes that share the same affordance motif. Within each such class the graphs are highly connected, since the affordance ratchet allows transitions to run in both directions. Between classes transitions are unidirectional, corresponding to a gain of affordance.

**Fig. 5:**
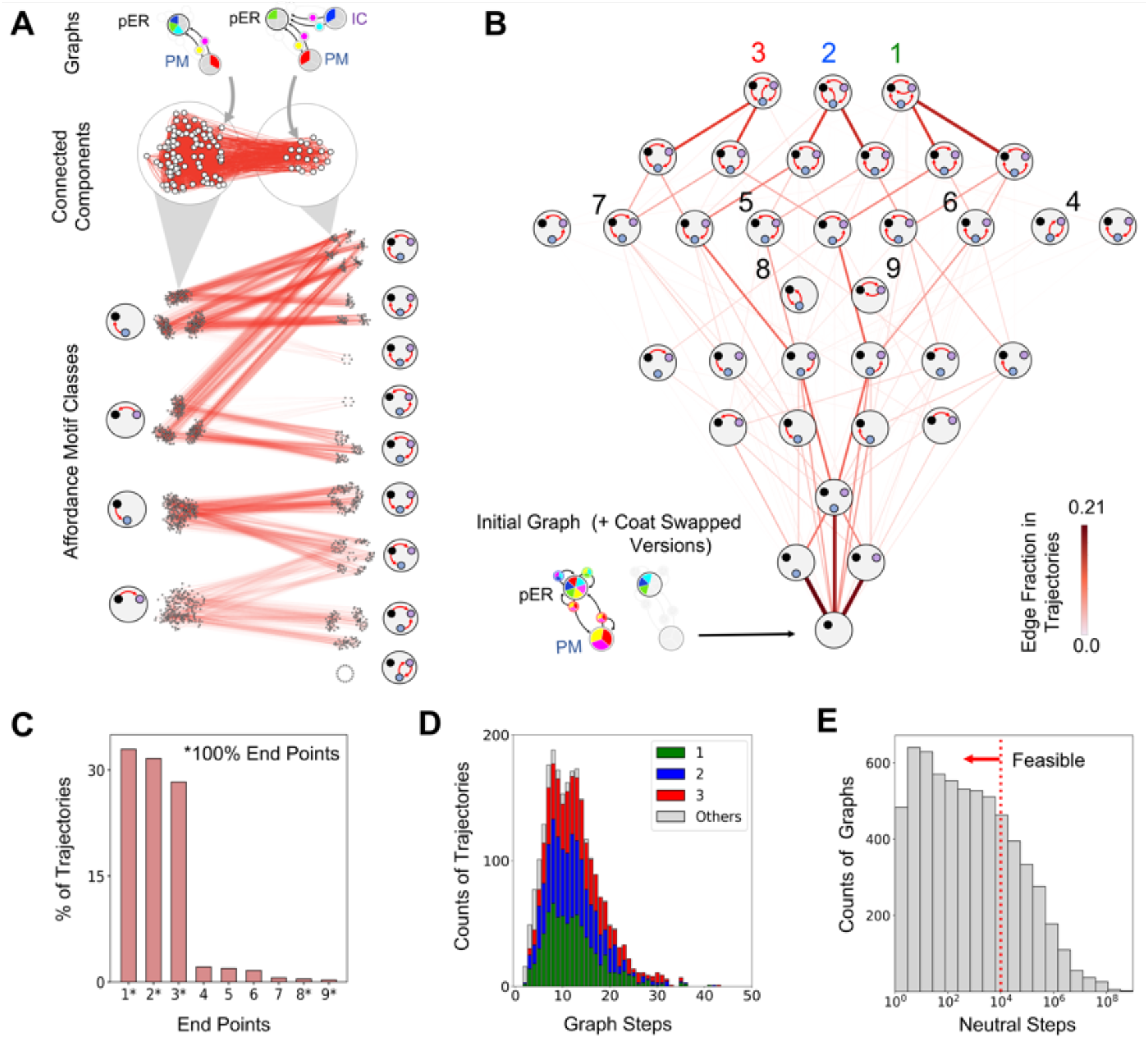
The evolutionary landscape and punctuated compartment gains. **(A)** Hierarchical structure of the evolutionary landscape. Individual compartment graphs are represented as nodes (white dots) connected by allowed transitions (directed red edges, corresponding to non-zero entries of the adjacency matrix). For the 3-coat activator model the evolutionary graph breaks into 216 connected components with bi-directional edges (clusters of white dots); each such component corresponds to a single affordance motif. Edges connecting one affordance motif class to another are unidirectional, corresponding to the expansion of the affordance motif. **(B)** The complete evolutionary landscape for the 3-coat activator model, containing 7,225 allowed 2-compartment and 3-compartment graphs. (The landscapes for other models are shown in Fig. S5.) Evolutionary trajectories are generated by a Monte Carlo simulation starting from the simplest proto-eukaryotic compartment graph (bottom-left) and randomly choosing any graph-to-graph transition allowed by the affordance ratchet, until no further transitions are possible. To represent this landscape we group individual graphs into affordance motif classes, as shown in (A). Edge thicknesses represent the fraction of simulated trajectories in which the corresponding transition type is observed. These classes are stratified vertically by increasing affordance complexity. The nine affordance motifs that contain endpoint graphs are numbered. **(C)** Distribution of endpoint affordance motifs. Approximately 80% of trajectories terminate at one of the three most complex endpoint motifs, a striking degree of canalization. Affordance endpoint classes 1, 2, 3, 8 and 9 contain only terminal graphs, while classes 4, 5, 6 and 7 contain both terminal and non-terminal graphs. **(D)** Distribution of trajectory lengths, defined as the number of distinct multigraphs sampled by a trajectory, inclusive of the starting and end points. Trajectory lengths have similar statistics across endpoint motifs 1,2, and 3, with a median value of 11. **(E)** Estimated lengths of neutral walks between successive graph transitions. Note the logarithmic scale of the x-axis. The dotted red line shows the cutoff number of steps that are feasible in the billion years separating FECA and LECA. See Fig. S5 for evolutionary landscapes under other SNARE regulatory models.

We represent a coarse-grained evolutionary landscape by treating affordance motif classes as nodes, each corresponding to a large number of individual graphs (Fig. 5B; Fig S5). These nodes connect to one another by edges whose weights represent how often the corresponding transitions are observed in sampled trajectories. This coarse-grained landscape is organized along two axes. Along the vertical axis, the number of labeled affordance features increases, representing stepwise gains in functional complexity. Along the horizontal axis, nodes contain the same number of labeled features but differ in which compartments or flows carry labels.

For the 3-coat activator model we simulate 3,000 trajectories on the 7,393 allowed graphs with 2 to 4 compartments (Fig. 5B). Since we are specifically interested in the emergence of the first IC compartment, we retain only the 2,437 trajectories that end in 3-compartment graphs while not traversing 4-compartment graphs. We find that approximately 80% of these 2,437 trajectories terminate at just three of the nine endpoint motifs (Fig. 5C), a striking degree of canalization. Certain endpoint motifs have no outward transitions, while others contain a combination of terminal graphs and outward transitions.

### Neutral walks, exaptation and punctuated evolution

The observed trajectories involve relatively few graph transitions, of order ∼10 (Fig. 5D). But this hides the complexity of the underlying dynamics. The first time some graph-*A* emerges along a trajectory, it does so with a specific molecular solution. This solution is almost never adjacent, in the sense of elementary evolutionary moves, to the solution of any graph-*B*. Starting from the modules present in the graph-*A* solution, gene duplications produce spare molecular copies which can randomly mutate, as long as these mutations still satisfy the graph-*A* conditions. This process moves along a neutral network [36, 37, 55], enabling exploration of the solution space. For a transition to happen, this neutral exploration must serendipitously end at a molecular solution of graph-*A* that is just one move away from a molecular solution of some graph-*B*. Though each step of the walk is not directly selected, the accumulated set of neutral changes is exapted to finally achieve the transition.

We can estimate the length of such a neutral walk statistically, under the simplifying assumption that successive steps are independent (Methods, Section J). Starting from some graph-*A*, we calculate the probability *P*_*AB*_ that a randomly sampled set of modules satisfies the graph condition of some neighbor graph-*B* at each step, while avoiding the graph-*A* forbidden region (entering which would be lethal). Each graph *A* can have several neighbors *B*. The neutral walk length will be geometrically distributed with a mean length1/∑_*B*_*P*_*AB*_. and graph-*B*^*\**^ will be chosen with probability *P*_*AB \**_ /∑ _*B*_*P*_*AB*_, The more neighbors graph-*A* has, and the easier the graph-*B* conditions are to satisfy, the shorter the neutral walk will be. There is an optimal number of duplicate copies for efficient neutral exploration: too many, and they enter the graph-*A* forbidden region; too few, and they do not satisfy the graph-*B* conditions. (Since the calculation of *P*_*AB*_ is an estimate, we chose to generate evolutionary trajectories assuming all neighbor graphs were equally likely.)

Applying the neutral walk estimation across all possible starting graphs, we find that the evolutionary dynamics are punctuated: periods of neutral exploration are separated by rare, sporadic jumps. The number of neutral steps between graph transitions is extremely broad, ranging from 10^0^ to 10^6^ (5E). Some neutral walks are short while others are exponentially longer, so the evolutionary trajectory is dominated by its longest step. This puts a cutoff on which transitions could feasibly occur in the billion years separating FECA and LECA. In the strong selection weak mutation limit with a per-gene per-generation mutation rate *µ*, neutral steps occur at a time separation of 1/*µ*. Assuming *µ* ∼ 10 ^−6^ typical of modern unicellular eukaryotes [56], and conservatively assuming a generation time of 30 days based on Asgard archaeal data [57], we see that ∼ 10^4^ steps are possible between FECA and LECA. 74% of neutral walks fall below this conservative limit, so most of the evolutionary landscape remains accessible.

## Discussion

Our study began with a puzzle: Molecularly allowed compartment graphs are extremely rare in the space of all possible graphs (Table I). How can evolution – driven only by gene duplication, deletion, and simple mutations (Fig. 2C) – discover complex endomembrane systems in such a large and sparse space?

The resolution of this puzzle comes from noticing that the molecular solutions corresponding to each graph allow for an extreme level of redundancy and neutrality. Since each compartment or vesicle typically contains multiple types of cargo, there are many ways of choosing appropriate budding and fusion modules. More importantly, the endomembrane system is spatially segregated by definition. This means certain combinations of cargo never co-occur on a given compartment, vesicle, or vesicle-compartment interface. Such non-occurring combinations enable the existence of a large number of neutral modules (Fig. 1F). Each allowed graph therefore has a combinatorially large collection of possible molecular solutions, all connected by elementary evolutionary moves.

The set of connected molecular solutions associated with a graph defines its neutral network [36, 37, 55]. If the neutral networks of two different graphs come within a single step of one another, then a graph transition becomes possible. Though allowed graphs are rare, their neutral networks are so large that each graph typically has hundreds of possible neighbors. As a result, we find that the entire evolutionary landscape forms a single connected component: it is possible to move between any pair of possible graphs by gene duplication, deletion and mutation (Fig. 5B). Each such transition requires a neutral walk [36, 37, 55] during which duplicate gene copies undergo mutations (Fig. 5E). The billion years separating FECA and LECA provides sufficient time for three quarters of molecularly allowed transitions to occur, and recombination could accelerate the process further [58]. This beautifully connects the molecular and geological timescales.

Within a connected landscape, entropy would favor trajectories that sampled the most highly connected regions. But these tend to be dominated by graphs with simple topologies. In order to move toward functional complexity, graph transitions must be biased by some selective pressure. To achieve this we introduce the idea of affordance labels (Fig. 3). These labels enable cellular functions to be localized to specific compartments and vesicle flows, and create a hierarchy of functional potential (Fig. 5B). Once we introduce the “affordance ratchet”, we find that evolutionary patterns tend to be highly reproducible. Molecular constraints exert a strong filtering effect on allowed graphs, and create highly contingent trajectories (Fig. 4). Almost all trajectories starting from a proto-eukaryotic initial state are canalized toward functionally complex endpoints (Fig. 6A,B, Fig. S6A). Across trajectories, affordance labels are acquired in a consistent order (Fig. 6C): the pER already begins with an affordance label; this is followed by the PM, then the IC; labeled flows come last. The first newly-added compartment is typically a subfunctionalized version of a pre-existing pER or PM, or neofunctionalized (Fig. 6D,E, Fig. S6B).

**Fig. 6:**
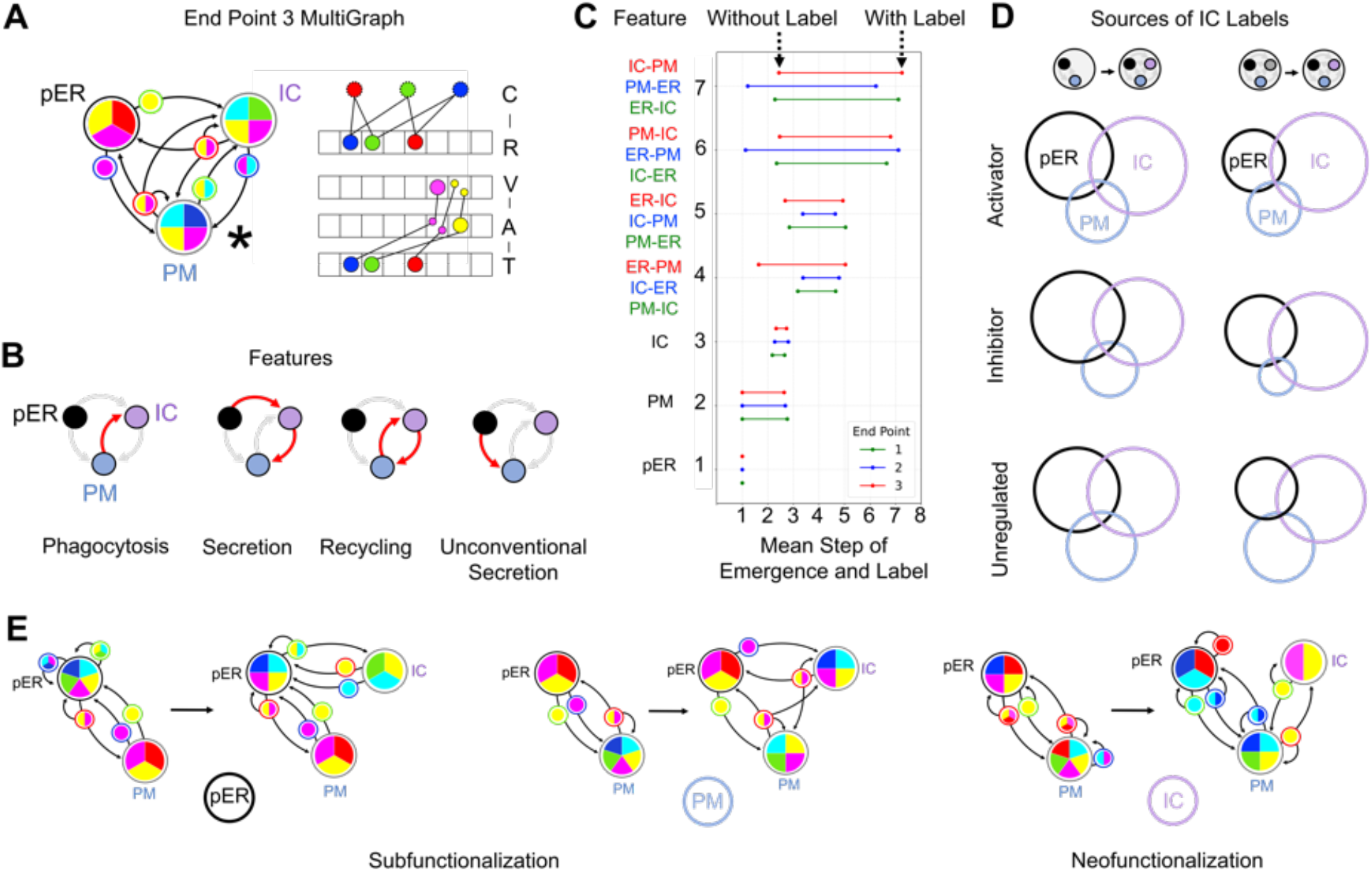
Emergence of novel compartments and functions. **(A)** An example graph from endpoint motif 3 for the SNARE activator model, and one of its molecular solutions (see Fig. S6A). **(B)** The graph shown in (A) facilitates biologically relevant vesicle flows coupling the pER, PM and IC. **(C)** The order in which compartments and flows acquire affordance labels. Among the 2,437 simulated trajectories of the SNARE activator model, we examine those terminating at endpoint motifs 1, 2 and 3 within at most 8 steps (inclusive of the initial and final). For each endpoint separately, we show the average step at which each indicated feature emerges and is subsequently never lost, along with the average step at which each feature first acquires an affordance label. All trajectories begin with an affordance-labeled pER and an unlabeled PM. The three endpoint motifs share the same early steps, then diverge in the order in which vesicle flows acquire affordance labels. **(D)** Function of the newly-added compartment. We analyze all transitions in which the IC first acquires an affordance label, across SNARE regulatory models. Left: transitions from 2-compartment graphs with a labeled pER and PM to 3-compartment graphs with a labeled IC. Right: transitions from 3-compartment graphs with a labeled pER and PM but an unlabeled IC to 3-compartment graphs with a labeled IC. We sample 10,000 transitions for each model, and determine how often the IC inherits an affordance label from the pER or the PM (subfunctionalization) or gains a new affordance label (neofunctionalization). The frequency of various cases is represented as an area-proportional Venn diagram. **(E)** Example of transitions in which a new IC inherits its label from the pER or PM (subfunctionalization), or gains a new label (neofunctionalization). See Fig. S6 for more statistics on the IC affordance label.

We see a fundamental link between molecular complexity and endomembrane complexity: having more coat types and more complex regulation expands the space of allowed graphs (Fig. S5). Systems with a single coat type produce only trivial graphs; at least three coat types, along with SNARE activation or inhibition, are required to support complex graphs (Table I). The emergence of multiple coat/adaptor complexes such as clathrin, COPI and COPII on the path from FECA to LECA [3, 34] may have been a prerequisite for endomembrane elaboration. Further, the existence of duplicate copies of coat recruiters, SNAREs and their regulators is essential, since this is what powers the neutral exploration of molecular space. However, the link between gene duplication and new compartments proposed by the organelle paralogy hypothesis is more nuanced than previously appreciated [33, 34]. There is no simple way to infer phenotype from genotype, nor phenotypic changes from genotypic changes. For every mutation that drives a graph transition, there are thousands of neutral mutations that leave no trace on the phenotype.

In our study we have attempted to go from mathematically possible, to molecularly allowed, to historically contingent, to actually observed endomembrane systems [59]. Each step in this hierarchy influences and restricts the possibilities of the next, culminating in the conserved cell plan of modern eukaryotes. The challenge going forward is to watch evolution in action, with experiments that explore how molecular constraints and selective pressures combine to drive endomembrane elaboration. Such experiments could directly test the predictions of our model: that compartment gains are preceded by periods of neutral molecular evolution, that early evolutionary choices constrain later ones, and that the function of newly-added compartments is shaped by subfunctionalization and neofunctionalization of pre-existing ones. This program will provide deep insights into the origins of eukaryotic cellular complexity, and explain the remarkable conservation and diversity of endomembrane systems across the eukaryotic tree of life.

## Resource Availability

See Supplemental Information for Figs. S1-S6 and Flowcharts 1-3. All code, simulation scripts, and data required to reproduce the results and figures presented in this manuscript are available in the GitHub repository at [Vesicle-Traffic-Model](https://github.com/sahanancbs/Punctuated-Evolution-of-Intracellular-Compartments-and-Vesicle-Traffic-in-Proto-Eukaryotes).

## Acknowledgements

MT acknowledges support from the Simons Foundation (287975). We thank Kabir Husain, Ishier Raote, Gautam Dey and Joel Dacks for their inputs, and all the members of the Simons Centre at NCBS for valuable discussions. This paper is dedicated to the memory of TTJ and TTV.

## Author Contributions

MT conceived the project. MT and SS developed the model. SS and KK carried out the analysis. MT and SS wrote the paper.

## Declaration of competing interests

The authors declare no competing interests.

## Methods

### A. Enumerating compartment graphs

We enumerate all possible directed multigraphs, where nodes represent compartments and directed edges represent vesicle flows. All edge multiplicities up to the maximal computationally feasible limits are included. We only include fully connected multigraphs, to account for membrane recycling. Topologically equivalent graphs are collapsed so that each distinct topology is represented exactly once. We assign coat labels to graph edges (Fig. 1B). Coat labels specify which cargo types can be loaded onto a vesicle via coat-cargo affinity.

All permutations of coat label assignments to edges are generated under the 2-coat and 3-coat models (larger coat sets are computationally infeasible). Multi-edges between compartments are retained only when they carry distinct coat labels. Finally, we label compartment identities (pER/PM for 2 compartments, pER/PM/IC for 3 compartments, pER/PM/IC1/IC2 for 4 compartments; Fig. 1C). Each fully labeled multigraph is a candidate compartment graph. The total number of graphs generated under each model configuration is shown in Table I.

### B. Determining compartment and vesicle compositions

Every cargo type is first synthesized at the pER. It then traverses the compartment graph by loading onto any vesicle whose coat it has an affinity for. To determine where a given cargo type ultimately accumulates, we construct a cargo-specific transport sub-graph (Fig. S1A). We collapse the strongly-connected components of the subgraph into nodes. These nodes then form a tree-like structure whose root is the pER and whose leaves are sinks [60, 61]. Each leaf is a terminal strongly-connected component. The cargo type will be found in high amounts on every compartment and vesicle belonging such a terminal component (Fig. S1A,B). The input of new cargo from the pER balances the degradation of cargo within each terminal component. We assume that the rate of degradation is much slower than the rate of transport. This means that any cargo type will only be present in trace amounts on its path from the pER to its terminal component, and need not be accounted for in determining the properties of the vesicles and compartments along this path.

### C. Budding and fusion modules

Certain cargo within the endomembrane system also serve regulatory roles. Each of these can be assigned a cargo type based on coat affinity (Fig. 1A). Compartments and vesicle compositions therefore specify the locations of regulatory molecules. Two molecular modules encode the core events of vesicle traffic in our model (Fig. 1D). The budding module, consisting of a coat (C) and its cognate recruiter (R); and the fusion module, consisting of a v-SNARE (V), its activator or inhibitor (A or I), and the cognate t-SNARE (T). We consider three models of SNARE regulation (SI Flowchart 1; Fig. S1E,F): (1) Activator, in which a v-SNARE is inactive except when co-localized with its activator on a vesicle. (2) Inhibitor, in which a v-SNARE is active except when colocalized with its inhibitor on a vesicle. (3) Unregulated, in which a v-SNARE is always active.

### D. Self-consistent molecular solutions

For a compartment graph with assigned compositions to be molecularly self-consistent, the following graph condition must hold. For each budding vesicle labeled with some coat, the source compartment must contain its cognate recruiter. For each fusing vesicle, the vesicle must carry the v-SNARE in an active state, and the target compartment must contain its cognate t-SNARE. To find a molecular solution for the SNARE activator model (and equivalently for other models), we search over all possible combinations of *n* × 2^*n*^ C-R doublets and 2^3*n*^ V-A-T triplets. We first eliminate any C-R doublets and V-A-T triplets that would enable any budding and fusion events absent from the graph. We then check if the remaining C-R doublets and V-A-T triplets collectively enable every budding and fusion event present in the graph (Fig. 1C,D). If this is successful, we have found a molecular solution. By successively removing C-R doublets or V-A-T triplets until failure, we can reach an irreducible solution. Each graph may have multiple irreducible solutions, reached by a different order of removal (Fig. S2A,B). We find the complete set of irreducible solutions by exhaustive search. This provides a compact representation of the solution space for each graph.

### E. Required, forbidden and neutral regions

The space of all possible modules can be partitioned into regions (Fig. 2A). Given a compartment graph with assigned compositions, we first define the forbidden region: all modules which enable a budding or fusion event which is absent from the graph. To meet the graph condition, no module can be in this region. Once these are eliminated, we group every module that enables each budding event, and group every module that enables each fusion event. Each such group defines a required region. The required regions can overlap, since the same module may enable multiple events. To meet the graph condition, there must be at least one module within each required region and none in the forbidden region. This defines a molecular solution. All remaining modules outside the forbidden and required regions comprise the neutral region.

### F. Elementary evolutionary moves

Three evolutionary moves act on the molecular solution of a compartment graph: duplication, deletion and mutation (Fig. 2C). A duplication adds a copy of an existing module, a deletion removes a module. (We assume modules are duplicated or deleted in their entirety.) A mutation flips a single coat-affinity bit of a cargo type in the module. The mutation move creates an adjacency structure in the space of all modules. A subset of the forbidden region, which we term unreachable, is only adjacent to other forbidden modules. Once the forbidden region is removed, we find that the remaining required and neutral regions form a single connected component for nearly every graph. (In 8.7% of 3-coat graphs the required/neutral C-R space breaks into disconnected components, but these can be connected via neutral spaces of other graphs.) This means that any pair of molecular solutions of a graph can be transformed into one another using elementary evolutionary moves that avoid the forbidden region.

### G. Graph transitions

We can check whether a given graph-1 can transition to a given graph-2 using elementary evolutionary moves. We first remove the union of the forbidden regions of graph-1 and graph-2 from the module space. We find that the remaining space always forms a single connected component under mutation (except in those rare cases discussed above, where the C-R space is disconnected). This means we do not have to explicitly construct the neutral walk from the given initial molecular solution to the transition step. If we start with a molecular solution of graph-1, it is always possible for gene duplications and mutations to create modules that prepopulate every required region of graph-2, as long as they do not enter the forbidden region of graph-1. It is therefore sufficient to look for one solution each of graph-1 and graph-2 that are a single move apart. The existence or non-existence of such a pair of solutions can be immediately inferred from the required and forbidden regions of the two graphs.

This procedure defines four types of graph transitions (Fig. 2G-J; SI Flowchart 2). (1) Bistable transition: the combined forbidden region does not fully contain any required region of either graph. A single solution can satisfy both graph conditions, and the initial state selects one of the graphs. (2) Mutation/Deletion transition: a single required region of graph-1 is fully within the forbidden region of graph-2. A deletion or any mutation into neutral space accomplishes the transition. (3) Mutation-only transition: a single required region of graph-1 is fully within the forbidden region of graph-2 and vice versa. If a mutation exists from the required region of graph-1 into the required region of graph-2, this accomplishes the transition. (4) Deletion-only transition: a single required region of graph-1 is fully within the unreachable forbidden region of graph-2. A deletion accomplishes the transition.

### H. Affordance labels and the affordance ratchet

Affordance labels quantify the functional potential of a compartment graph. A compartment has an affordance label if at least one cargo type resides exclusively on that compartment, providing a unique molecular identity marker. A vesicle flow between labeled compartments has an affordance label if some cargo type from the source compartment flows exclusively to the target compartment (SI Flowchart 3). The pER always carries an affordance label because cargo type 0 never leaves the pER. The set of labeled compartments and flows defines a graph’s affordance motif (Fig. 3A).

Consider any pair of graphs between which a transition is possible using elementary evolutionary moves. Under the affordance ratchet, the transition must preserve the affordance motif or expand it (Fig. 3B). Importantly, to preserve a labeled compartment or vesicle flow, the set of labels before and after the transition must overlap (Fig. 3C). If the graphs share an affordance motif then the transition is allowed in both directions. If one graph’s affordance motif is a subset of the other’s, then the transition is allowed from the first to the second. In all other cases the transition is not allowed.

### I. Evolutionary trajectories

A trajectory is a sequence of graph transitions. To construct trajectories, we first list all allowed graphs under a given coat/SNARE model. We next list all directed transitions allowed by elementary evolutionary moves and the affordance ratchet. We store this information as a binary adjacency matrix. We then perform a Monte Carlo simulation, starting from the proto-eukaryotic initial graph in all its coat-swapping permutations (Fig. 5B, bottom). At each step, we randomly select and jump to one of the outward neighbors of the current graph. This continues until we reach an endpoint graph, one with no outward neighbors.

### J. Neutral walk lengths

Suppose we are given some graph-*A* and graph-*B*, between which we have established that a transition is possible. We start with some molecular solution of graph-*A*. Now we populate the genome with spare modules through the duplication of existing modules. The neutral walk driven by the mutation of these spare modules can be approximated by assuming that, at each step, the entire set of modules is re-sampled at random with probability *ρ*. The real neutral walk is slower, since only one module is mutated at a time. The neutral walk must ensure that there is always at least one module in each required region of graph-*A*, and none in its forbidden region. For simplicity we focus only on the second condition: a module entering the graph-*A* forbidden region is lethal.

Now we ask how this neutral walk can take us within one step of a graph-*B* solution. The graph-*B* condition defines several overlapping required regions. We consider this as a Venn diagram, and separate it into non-overlapping zones. A minimal subset of zones that hits every required region is called an irreducible cover. To satisfy the graph condition, each zone in the cover must contain at least one module, and the forbidden region must contain none. Each graph typically has many possible irreducible covers. Let *y*_*A*_ be the size of the graph-*A* forbidden region. Remove this region from the space of possible modules, to get a reduced space. Let *y*_*B\A*_ be the size of the graph-*B* forbidden region within the reduced space. Now consider some irreducible cover of graph-*B* with several zones, each of size *z*_*i*_, *i* = 1,... in the reduced space. Cycle over covers until we get one where either all the *z*_*i*_ are non-zero (bistable transition) or all-but-one of the *z*_*i*_ are non-zero (mutation/deletion transition). Let *k* be the number of non-zero *z*_*i*_. The probability that modules sampled with probability *ρ* satisfy the graph-*B* condition while avoiding the graph-*A* forbidden zone is:

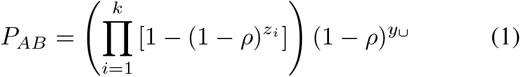

where we have set *y*_*⋃*_ = *y*_*B\A*_ + *y*_*A*_ as the size of the union of forbidden regions. We ignore the requirement that there should be no graph-*A* module in the graph-*B* forbidden region, as this can easily be removed by deletion.

For the 3-coat activator model we typically have *k ∼* 5 zones in the cover, each of size *z*_*i*_ *∼* 5 (Fig. S2A,B); and the forbidden region is of size *y*_*⋃*_ *∼* 200 (Fig. S2C,D). Under these conditions the *ρ* which maximizes *P* will be small, so Eq. (1) becomes

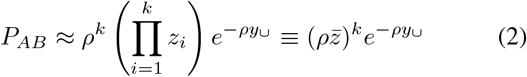

where 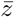 is the geometric mean of the non-zero zone sizes. *P* is maximized at ρ^*^ = (*k/y* _*∪*_). If the space of possible modules is of size *N*, then the optimal number of paralogs is *N ρ* ^*\**^. Substituting this into Eq. (2),

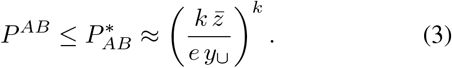

The total probability of reaching some neighbor of graph-*A* at each step can be upper-bounded by maximizing each term in the sum below independently with respect to *ρ*:

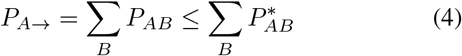

The mean number of neutral steps until a transition occurs, including lethal steps, is lower-bounded by the inverse of this probability. The total number of non-lethal steps will be lower-bounded by

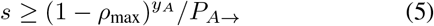

where *ρ*_max_ is the largest *ρ* found while maximizing the individual *P*_*AB*_. We compute this quantity for each graph-*A*, and plot its distribution in Fig. 5E as an estimate of the neutral walk length.

## Supplemental Information for

### Supplemental Figures AND Flowcharts

**Fig. S1:**
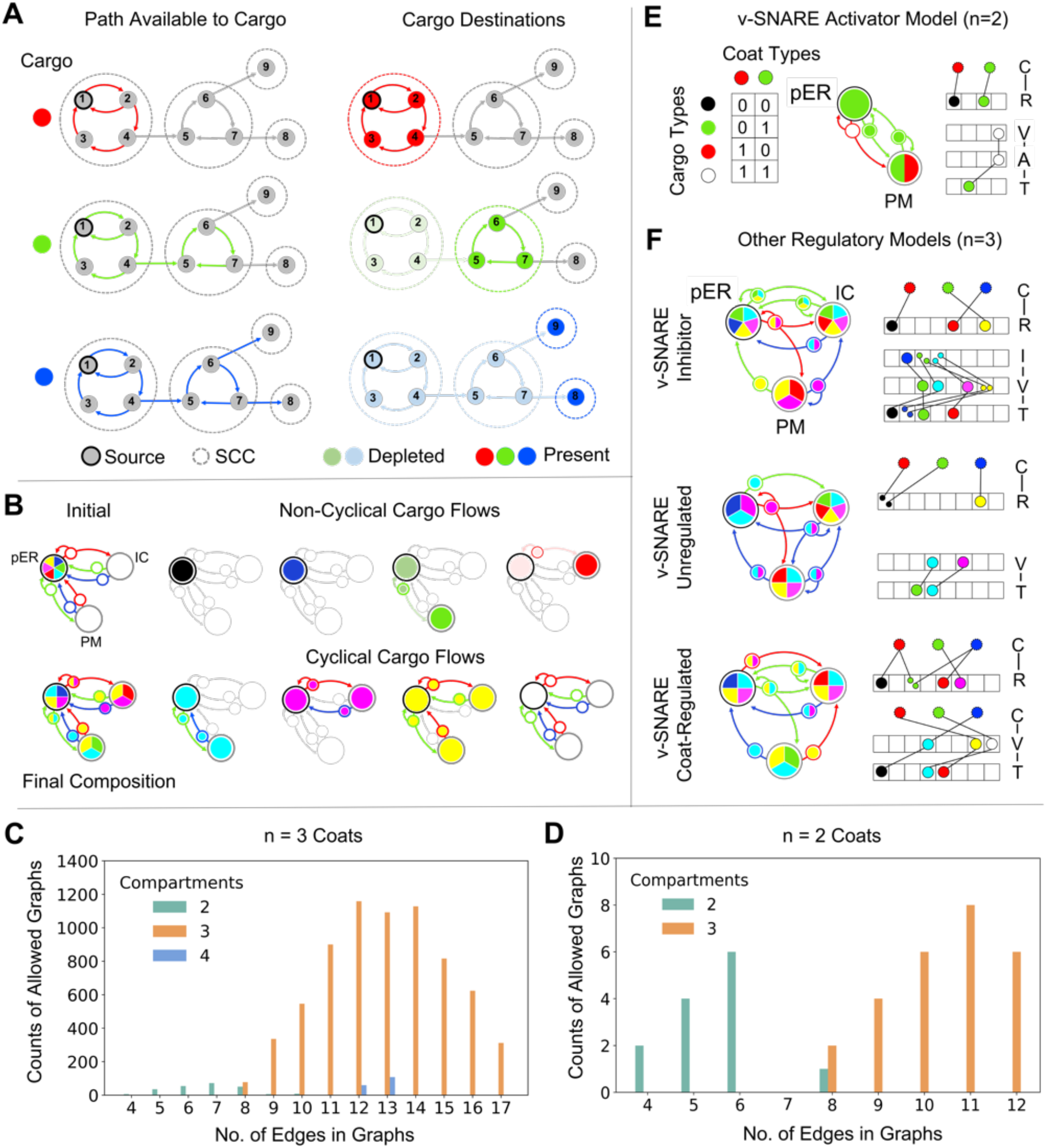
Cargo flows, compartment compositions, and SNARE regulation. **(A)** All cargo is synthesized at the pER and follows the vesicle flows accessible to it. For each cargo type we construct the transport subgraph, formed by edges with coat labels compatible with that cargo type. We condense this subgraph into a tree of strongly connected components rooted at the pER. Cargo accumulates on all the compartments and vesicles of terminal components, but are depleted on the path from the pER to that component. **(B)** In 3-compartment graphs, cargo types with non-cyclical flows are present only on compartments. Cargo types with cyclical flows are present on compartments and vesicles. **(C**,**D)** The distribution of compartment and edge numbers across allowed graphs with coat-labeled edges and a pER-identified node. **(E**,**F)** Examples of allowed graphs under different SNARE regulatory models: Activator, Inhibitor, Unregulated, and Coat-regulated. Our analysis is restricted to the first three models.

**Fig. S2:**
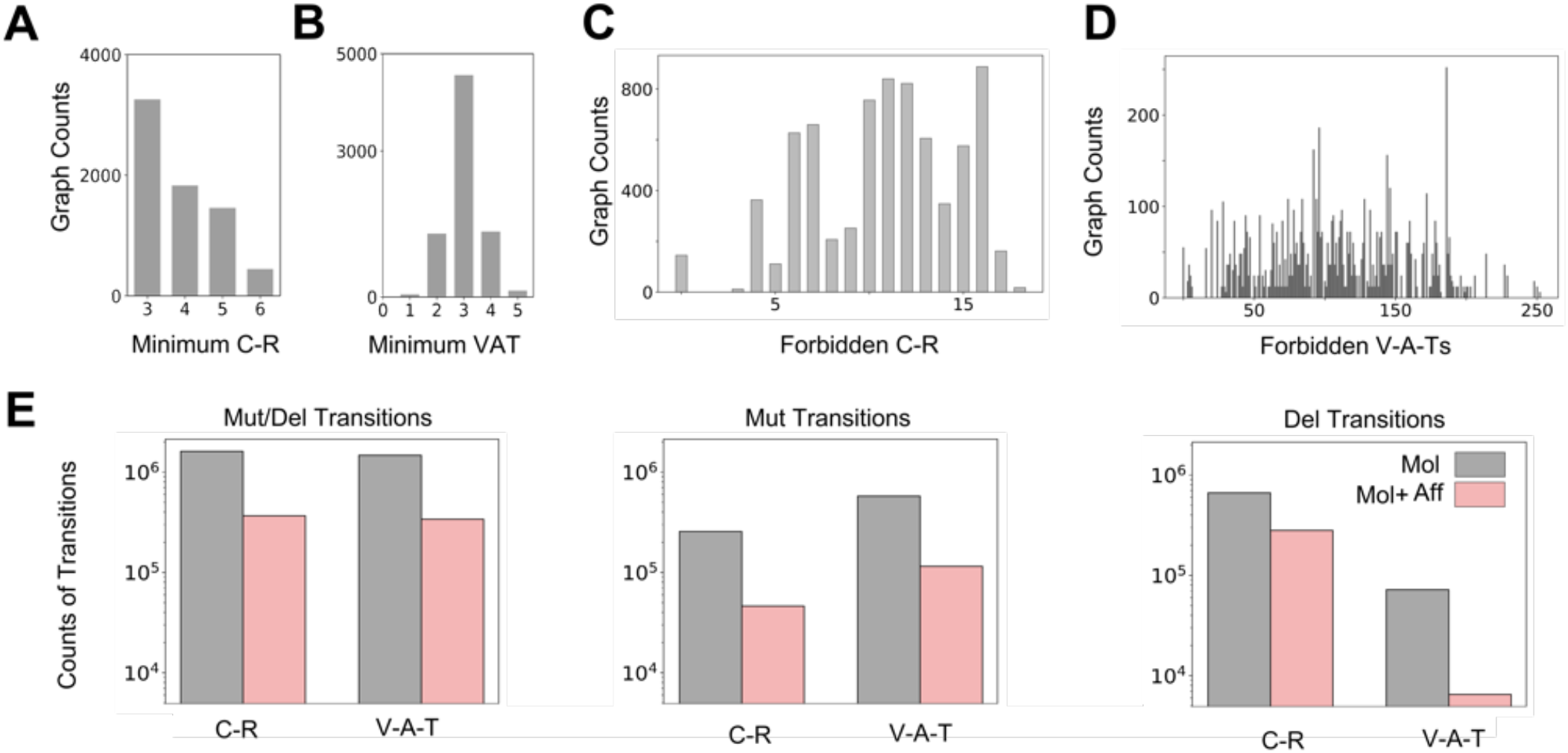
Required modules, forbidden modules, and graph transitions. **(A**,**B)** The distribution of the smallest set of (A) C-R modules and (B) V-A-T modules in molecular solutions, across all allowed graphs. **(C**,**D)** The distribution of the size of the forbidden region for (C) C-R modules and (D) V-A-T modules. **(E)** The number of graph transitions, classified by the type of elementary evolutionary move driving the transition. Transitions are further grouped by whether the move involves C-R modules or V-A-T modules. We compare all graph transitions against the subset permitted under the affordance ratchet.

**Fig. S3:**
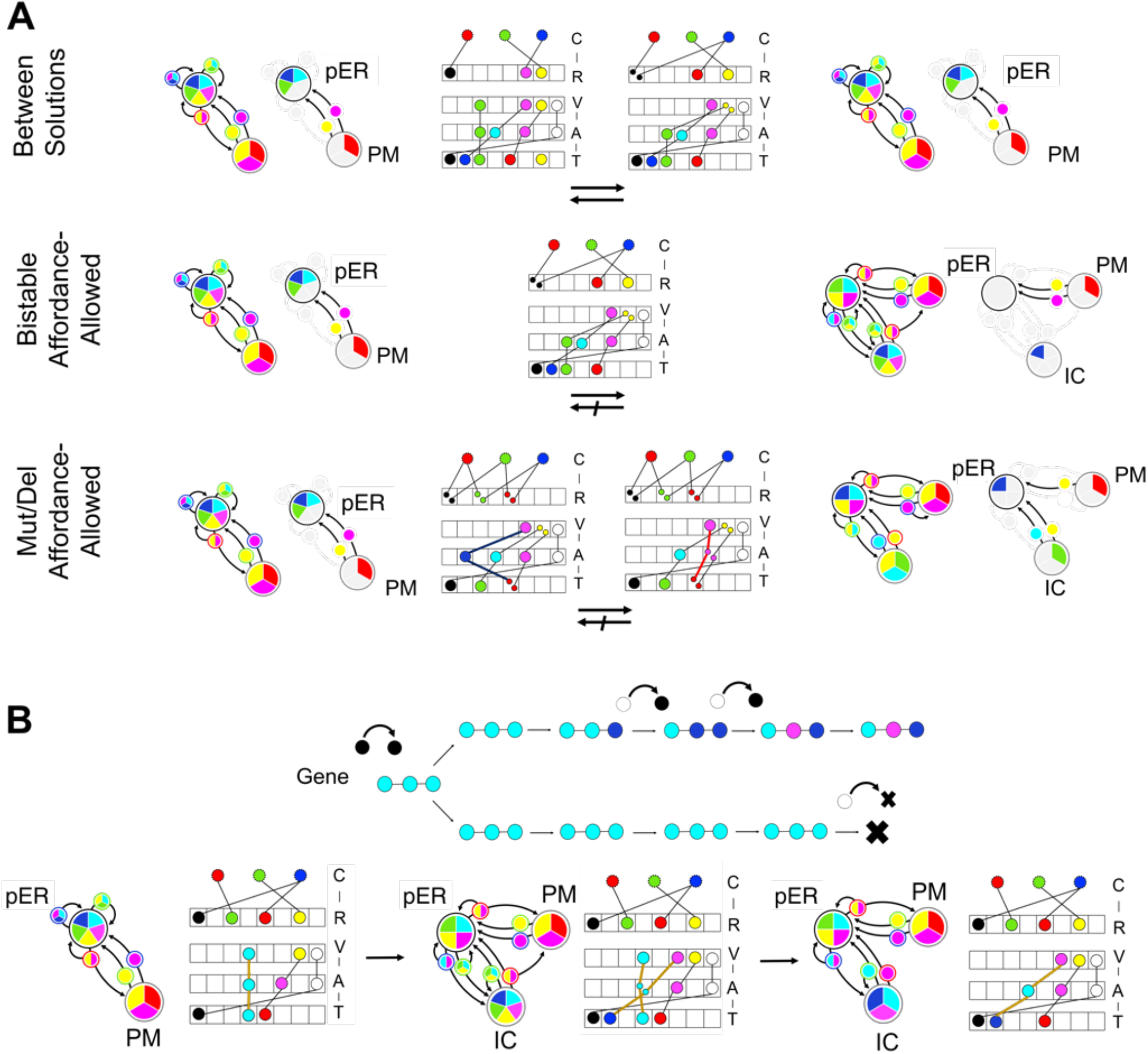
Evolutionary accessibility of molecular solutions and graph transitions. **(A)** Transitions between molecular solutions of initial and final graphs. The first row shows a case where the same graph has multiple solutions. The second row shows a case where the same solution is compatible with multiple graphs. The third row shows a mutation-driven transition. **(B)** Explicit example of how gene duplication, mutation and deletion achieve transitions between different molecular solutions of successive graphs.

**Fig. S4:**
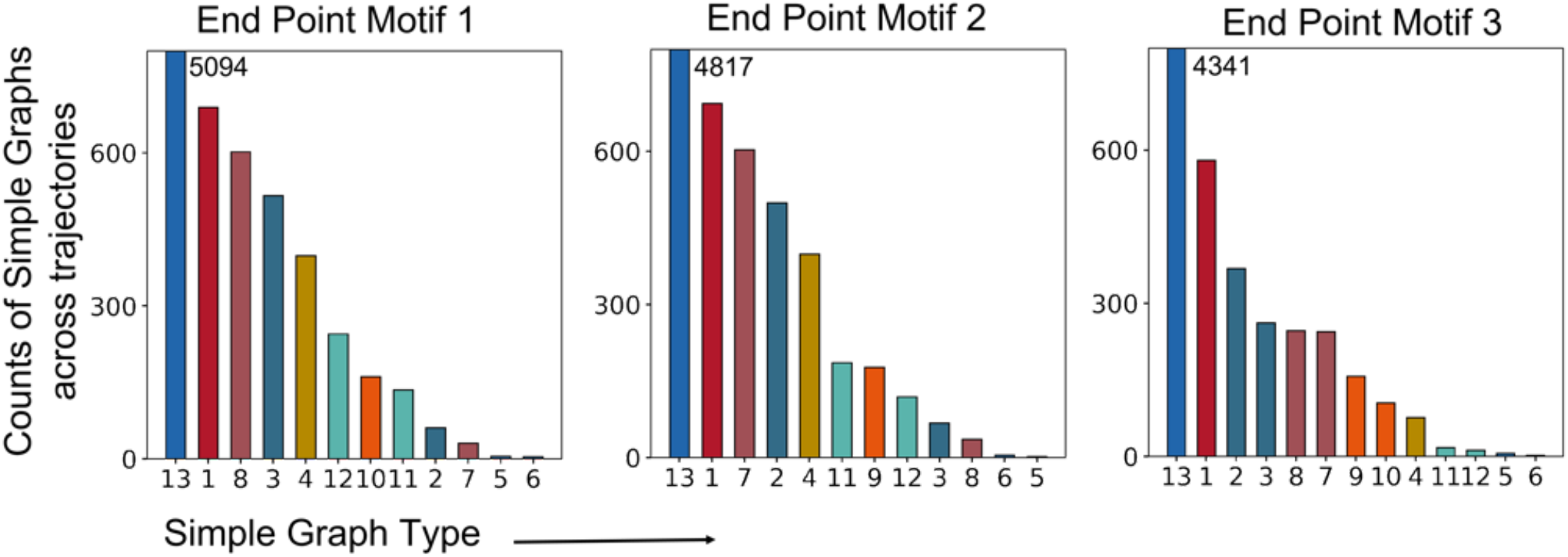
Endpoint-specific biases and symmetry effects in evolutionary trajectories. The 2,437 trajectories simulated under the 3-coat activator model are grouped into nine classes based on their endpoint affordance motif. For each group, all simple graphs visited along the trajectories are pooled and the frequency of each simple graph is computed within that group. We show frequency distributions for the three most common endpoint motifs. These distributions reveal several features of evolutionary contingency. First, certain simple graphs are over-represented across trajectories terminating at all endpoint motifs, reflecting topological symmetries that make them broadly accessible. Simple graph type 13 is an example of this. Second, some simple graphs preferentially lead to specific endpoints, and early choices constrain later possibilities. For example, trajectories passing through simple graph type 11 rarely terminate at endpoint motif 3. Third, symmetry pairs of simple graphs, obtained by exchanging PM and IC labels relative to the pER, show complementary endpoint preferences. Within such pairs, one member is over-represented in trajectories terminating at a given motif, while its partner is under-represented in that motif but favored in the mirrored endpoint. When the endpoint motif itself is symmetric, both members of a symmetry pair are represented at comparable frequencies. For example, the symmetry pair 7-8 shows biased representation toward motifs 1 and 2 respectively, but near-equal representation in motif 3.

**Fig. S5:**
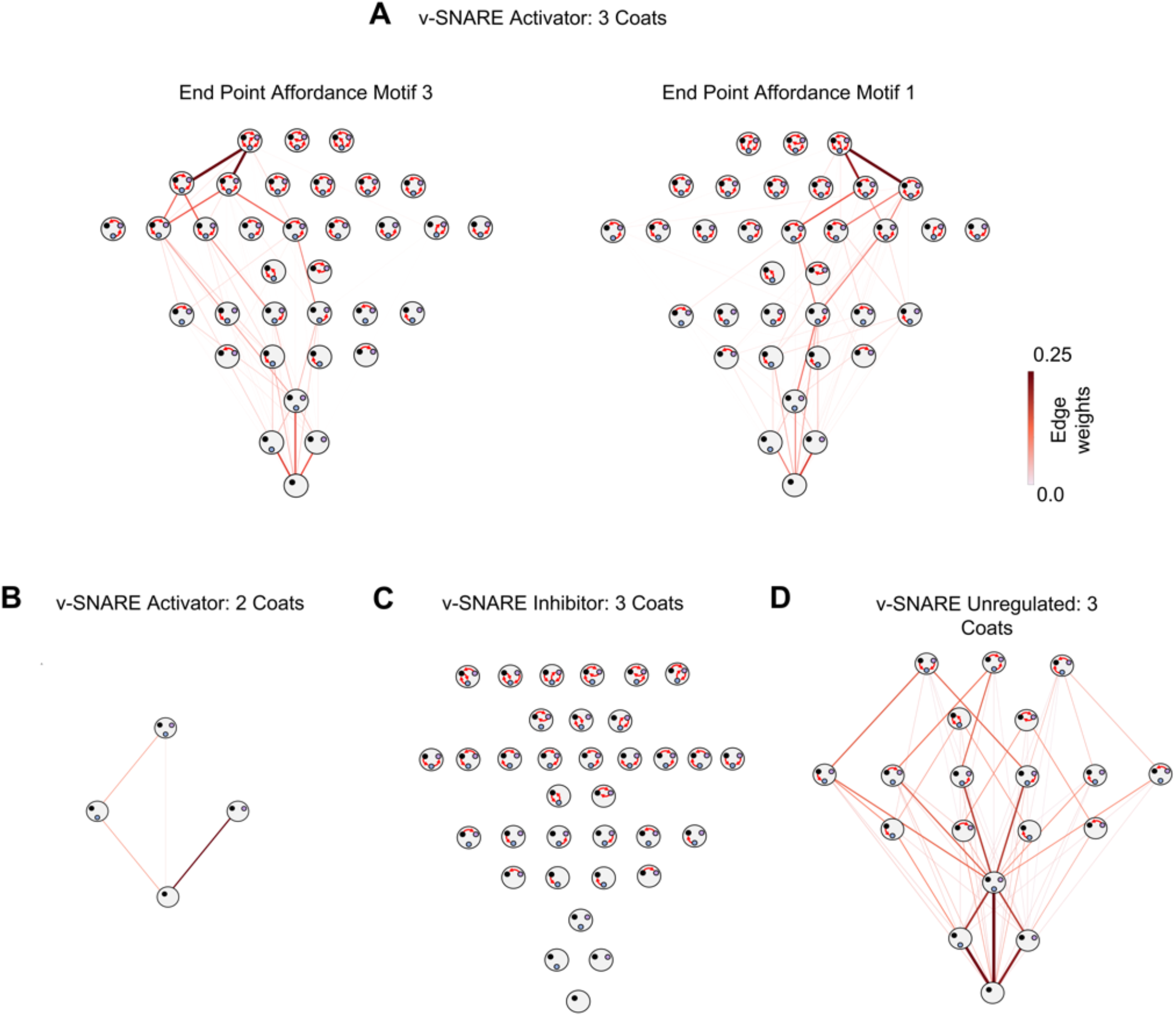
Evolutionary landscapes across coat type number and SNARE regulatory models. **(A)** Evolutionary landscapes for trajectories terminating at motifs 1 (803 of 2437 simulated trajectories) and 3 (690 of 2437 trajectories) of the v-SNARE Activator model, shown separately. Motif 1 accumulates more terminating trajectories because there are more accessible evolutionary routes leading to it, with trajectories exploring multiple lower-complexity layers before arriving at the endpoint. **(B)** Evolutionary landscape of the 2-coat model, showing 1000 simulated trajectories. **(C)** Evolutionary landscape of the SNARE inhibitor model. The complexity of affordance motifs is comparable to that of the activator model. Certain motifs absent in the activator model are present here, demonstrating that different SNARE regulatory models give rise to different accessible graph spaces. We do not show trajectories for the inhibitor model. This model has a very large number of graphs and transitions, and it is not feasible to determine its complete adjacency motif. Trajectories must be generated by sampling, which is highly inefficient. **(D)** Evolutionary landscape of the unregulated SNARE model, showing 1938 of 2000 simulated trajectories that end in 3-compartment graphs and do not traverse 4-compartment graphs.

**Fig. S6:**
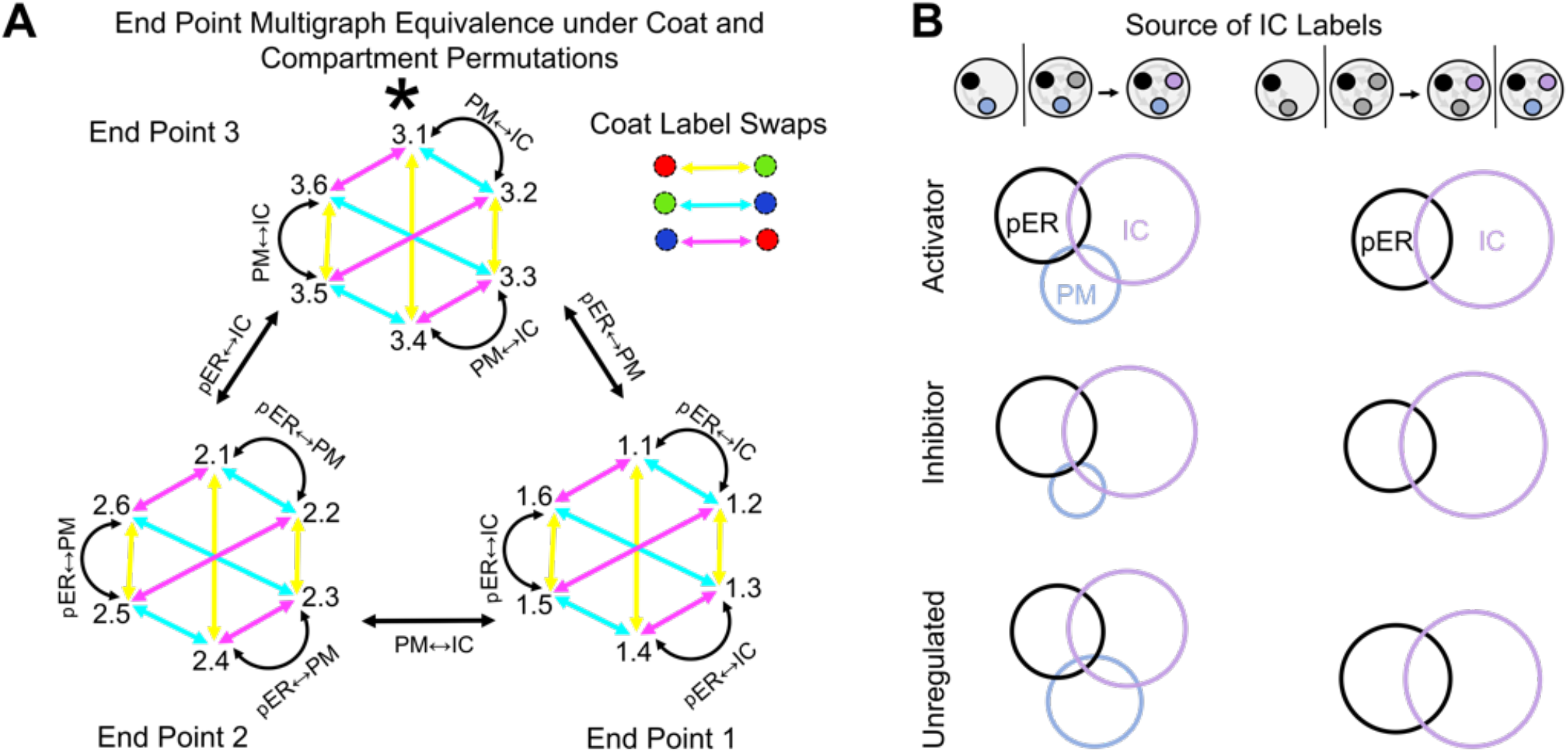
Canalization of the evolutionary landscape and sources of IC labels with labeled or unlabeled PM. **(A)** Endpoint motifs 1, 2 and 3 of the v-SNARE Activator model each contain only a single compartment-graph type once compartment identity (pER/PM/IC) and coat-label (red/green/blue) permutations are accounted for. The * graph is illustrated in Fig. 6A. This demonstrates that the evolutionary landscape is highly canalized due to molecular constraints and the action of the affordance ratchet. **(B)** Sources of IC affordance labels. For each v-SNARE model (activator, inhibitor, and unregulated), we sample random pairs of allowed graphs in which the first graph either has no third compartment or has an unlabeled third compartment, and the subsequent graph contains a labeled third compartment (IC). We retain only transitions that are molecularly allowed and affordance-preserving or affordance-expanding, until 10,000 such valid transitions are obtained. These transitions are then classified into two cases. Left: the first graph has a labeled PM. Right: the first graph has an unlabeled PM. Area-proportional Venn diagrams for these two cases, across the three SNARE regulatory models, summarize the inferred origins of IC labels: purely from the pER, purely from the PM, novel at the IC, or mixed cases.

**Flowchart 1.**
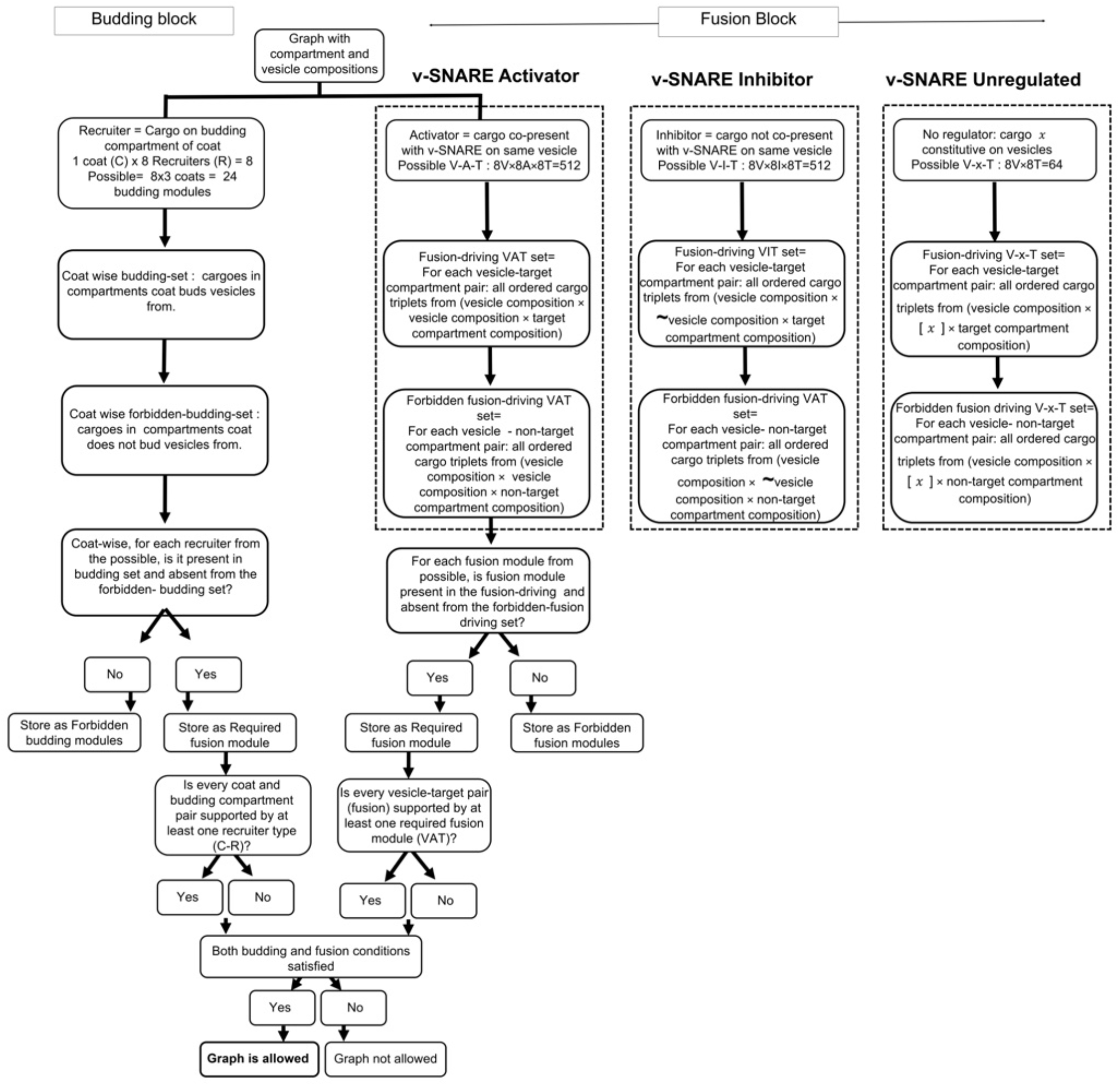
SNARE regulatory models. The three regulatory models considered in this paper (activator, inhibitor, and unregulated) specify the conditions under which a v-SNARE becomes active on a vesicle. Each model imposes different constraints on which molecular configurations satisfy the fusion condition of a compartment graph.

**Flowchart 2.**
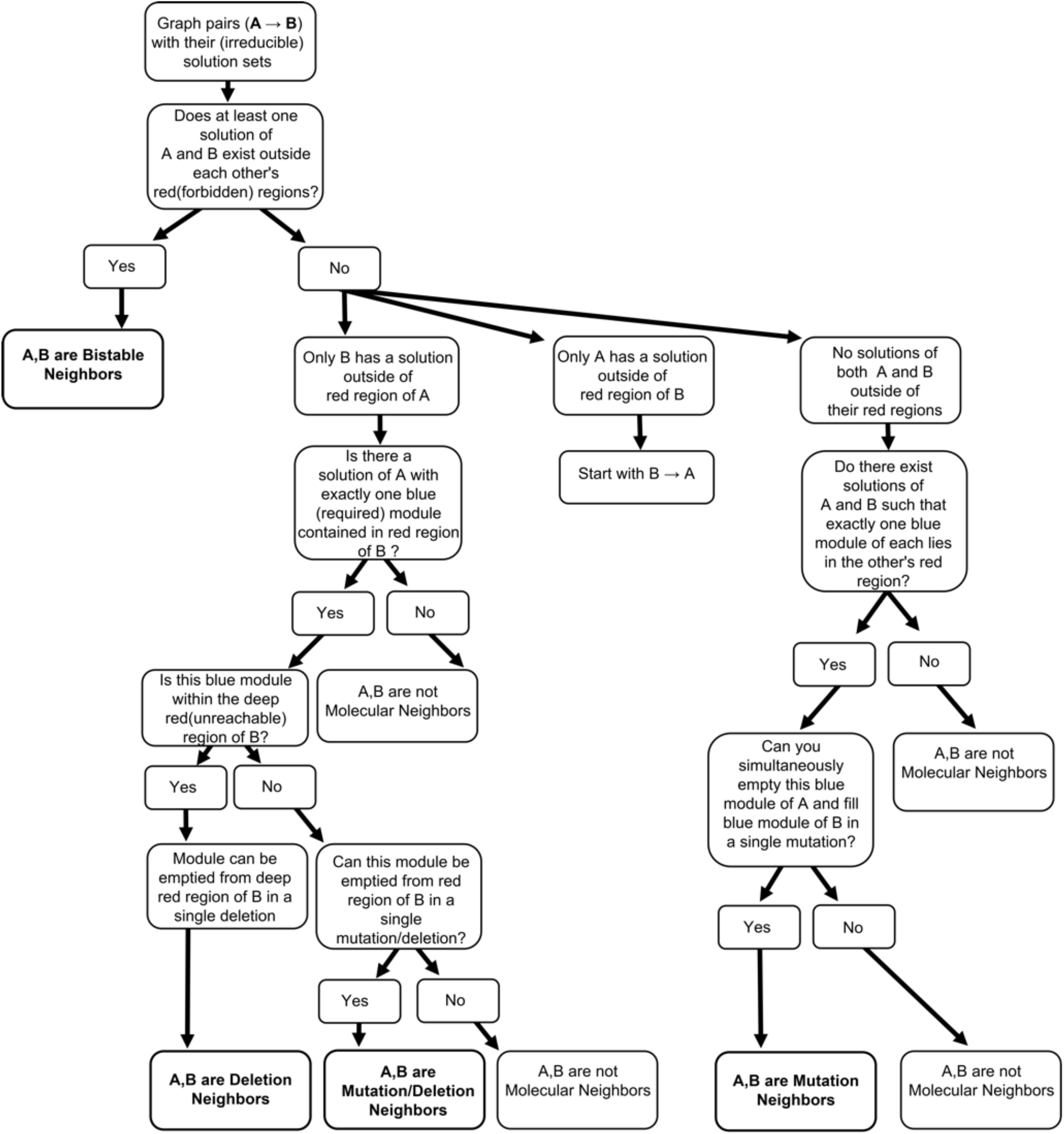
Graph transitions: bistable, mutation/deletion, mutation-only and deletion-only. For each pair of allowed compartment graphs, this flowchart specifies the procedure for determining whether a graph transition exists and classifying it by the nature of the underlying elementary evolutionary move.

**Flowchart 3.**
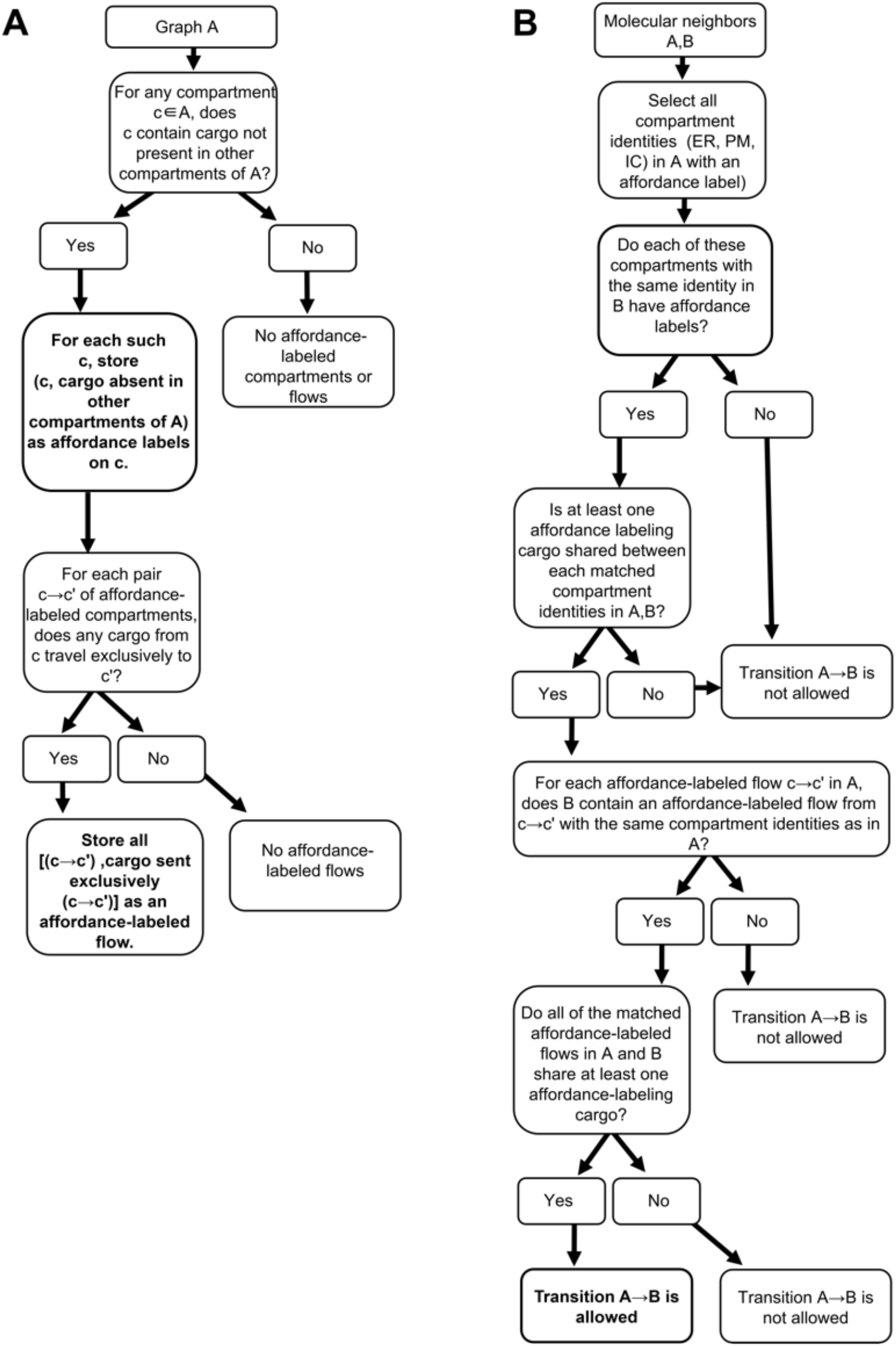
Affordance labels and the affordance ratchet. **A** The procedure for assigning affordance labels to compartments and vesicle flows of a given compartment graph. **B** The affordance ratchet: the rule by which graph transitions are classified as selected against, neutral, or representing an evolutionary leap, based on whether they destroy, preserve, or expand the affordance motif.

## Notes

https://github.com/sahanancbs/Punctuated-Evolution-of-Intracellular-Compartments-and-Vesicle-Traffic-in-Proto-Eukaryotes

### Competing Interest Statement

The authors have declared no competing interest.

### Summary of Updates

We have added a constraint on the lengths of neutral walks. This is based on the analysis of the number of mutations that are feasible in the billion year period separating FECA and LECA. Fig. 5E has been revised to reflect this cutoff on neutral walk lengths. The text has been modified to reflect the fact that neutral walk lengths cannot be unbounded.

